# An Inositol Receptor Orchestrates Carbon Utilization and Fungal Virulence

**DOI:** 10.64898/2026.03.02.708983

**Authors:** Yina Wang, Robert Tancer, Maggie Wear, Katrina Jackson, Yu Zhang, Yong-Gui Gao, Kirsten Nielsen, Arturo Casadevall, Chaoyang Xue

## Abstract

Nutrient sensing and utilization are critical for microbes to adapt to their environment and evade predators. The encapsuled yeast *Cryptococcus neoformans* has a unique ability to sense and utilize the sugar inositol as both a signaling molecule, and as a carbon source. This trait is advantageous not only in the natural environment, but to promote its pathogenesis, including invasion of the central nervous system. Unlike many other fungal species, which utilize the canonical carbon catabolite repression (CCR) mechanism to prioritize glucose metabolism, we have identified a novel inositol transporter-like receptor (transceptor) Itr4 that also regulates CCR genes to maintain inositol metabolism activity even under high glucose conditions. Itr4 binds inositol and regulates the function of other inositol transporters (*ITR*s) and its loss leads to a significant decrease in inositol uptake activity, resulting in a lack of mating, reduced capsule size, reduced blood brain barrier (BBB) penetration, and virulence attenuation. Mutagenesis analysis of Itr4 protein identified an essential N-terminal tail and two amino acids, Q388 and Q389, as sites of functional importance. The *ITR4^Q388A, Q389A^* allele showed a dominant phenotype with increased inositol uptake, enlarged capsule and significantly attenuated virulence. In summary, we identify a new *C. neoformans* inositol transceptor, Itr4, the first such example in eukaryotes, with novel regulatory roles in both inositol and glucose metabolism and fungal virulence.

## Introduction

The human fungal pathogen *Cryptococcus neoformans* is the leading cause of life-threatening fungal meningoencephalitis that accounts for ∼15% of AIDS-related deaths annually[1, 2]. Cryptococcal meningitis occurs after aerosolized fungal cells or spores dispersed in the environment are inhaled and disseminate from the lung to the brain via the blood stream[3, 4]. The medical significance and genetic tractability of *C. neoformans* have propelled extensive studies on the mechanism of its virulence with significant progresses in identifying factors required for fungal pathogenesis[5–8],. However, our understanding of the molecular basis of cryptococcal pathogenesis, including meningoencephalitis remains incomplete.

In previous studies we have shown the importance of fungal myo-inositol acquisition and metabolism in *Cryptococcus* species[9]. Interestingly, environmental reservoirs for *Cryptococcus*, such as *Eucalyptus spp*, which is also important for cryptococcal mating, as well as the natural fungal predator amoeba both contain high levels of inositol[9, 10]. In addition to environmental niches, inositol is highly abundant in the brain, promotes fungal traversal of the blood brain barrier (BBB) and plays a critical role in host-pathogen interactions during infection of the central nervous system (CNS)[11–13]. The cryptococcal genome also reflects the evolutionary adaptations associated with the expanded role of inositol in this organism[9, 11]. Therefore, we reasoned that an evolutionary specialization that may have originally promoted survival on natural hosts also facilitates the infection of the human host.

Fungal cells acquire inositol through two mechanisms. One is the internal synthesis of inositol. Intracellular glucose can be converted into inositol in a multiple-step biosynthetic pathway[14, 15]. Inositol can also be imported from the extracellular environment via inositol transporters[16–20]. The inositol transporter (*ITR*) gene family is part of the sugar transporter superfamily and plays an important role in inositol acquisition in fungi. Our previous studies showed that *C. neoformans* contains an expanded *ITR* gene family in its genome[11]. The *C. neoformans ITR* homologues can be divided into two distinct subgroups (I and II). The seven members of subgroup I (*ITR1, ITR1A, ITR2, ITR3, ITR3A, ITR3B,* and *ITR3C*) share high sequence homology with well-characterized *ITR*s in other fungi, such as *S. cerevisiae* and *C. albicans*. Subgroup II has three members (*ITR4, ITR5,* and *ITR6*) that are more distantly related *ITR*s that are more closely related to *HGT19*, a potential second *ITR* in *C. albicans*[21]. The expansion of this gene family suggests an important biological significance for inositol sensing, transporting and utilization in this fungus. Our previous functional studies on the seven group I *ITR*s have revealed that Itr1A and Itr3C are major low-affinity inositol transporters required for fungal sexual reproduction and fungal virulence in murine infection models[21]. The *itr1a*Δ *itr3c*Δ double mutant has a virulence defect, with a reduced ability to penetrate across the BBB that leads to virulence attenuation during brain infection[13]. This result suggests that inositol acquisition is important for the pathogenicity of *C. neoformans*. Because fungal *ITR*s are proton-dependent importers, distinct pharmacokinetically from sodium-dependent human and animal myo-inositol transporters (MITS)[20], fungal *ITR*s may be developed as a valuable drug target.

Genome wide association studies (GWAS) conducted on clinical isolates collected during the Cryptococcal Optimal Antiretroviral Therapy Timing (COAT) Trial in Uganda[22], also identified *ITR4,* a member of subgroup II, as one of several genes having polymorphisms associated with differential human immune responses and patient survival[23, 24], suggesting the clinical importance of inositol utilization. However, the mechanisms for how fungal cells sense, take up and utilize intracellular inositol to influence its development and virulence remain poorly understood. The function of subgroup II *ITR*s (Itr4, Itr5 and Itr6) has not been characterized.

In this study, we analyzed the function of the group II *ITR* genes in *C. neoformans* and found that Itr4 was the only one with significant impact on mating and fungal virulence. Using a *S. cerevisiae* heterologous expression system, we confirmed that Itr4 functions as an inositol sensor rather than a transporter. Mutagenesis studies showed that Itr4 had an outsized role in regulating fungal inositol uptake and fungal virulence in a murine model of systemic cryptococcosis. We further identified the potential inositol binding sites in Itr4 and generated *ITR4^Q388A, Q389A^* mutant allele based on site mutagenesis. Significantly, although the *itr4*Δ mutant produced a smaller capsule while the *ITR4^Q388A, Q389A^* allele produced an enlarged capsule, both mutants showed significant virulence attenuation in murine infection models. These results support the conclusion that inositol sensing, uptake and utilization in *C. neoformans* is important for the *Cryptococcus*-host interaction, BBB crossing and fungal virulence.

## Results

### Itr4 is a group II *ITR* homolog required for polysaccharide capsule formation

The production of a polysaccharide capsule is a key virulence factor that plays a critical role in *Cryptococcus* pathogenesis. Previously we demonstrated that inositol regulates both capsule size and polysaccharide structure[21, 25]. We also identified the expanded gene family of inositol transporters (*ITR*s) and inositol oxygenase (*MIO*s) and revealed that both two major group I *ITR*s (*ITR1A* and *ITR3C*) and inositol catabolic pathway (*MIO*s) are required for fungal virulence[25]. Despite a clear virulence attenuation of the *itr1aΔ itr3cΔ* double mutant, the individual mutants of group I *ITR*s did not have a significant defect on capsule production, likely due to functional redundancy. In addition, the function of the group II *ITR*s (*ITR4, ITR5* and *ITR6*) in *C. neoformans* has not been studied in detail. To further understand the role of the group II *ITR*s in capsule regulation, we generated the *itr4Δ, itr5Δ,* and *itr6Δ* single mutants and their triple mutants (*itr4Δ itr5Δ itr6Δ*) and measured the size of their capsules. We found that while the *itr5Δ* and *itr6Δ* single mutants did not exhibit obvious capsule difference, both the *itr4Δ* single mutant and the triple mutant cells had significantly reduced capsule and cell size (Fig 1A and 1B). The capsule defect in the *itr4Δ* mutant background was fully complemented by reintroducing the *ITR4* allele, indicating that Itr4 is required for capsule growth under this condition. In addition to having a capsule, thermotolerance and melanin production are two other classical virulence factors in *C. neoformans*. Our phenotypic analysis showed that all three *itr* single mutants and the triple mutants had normal growth at 37°C with proper melanin production. (S1A and S1B Figs).

**Figure 1.**
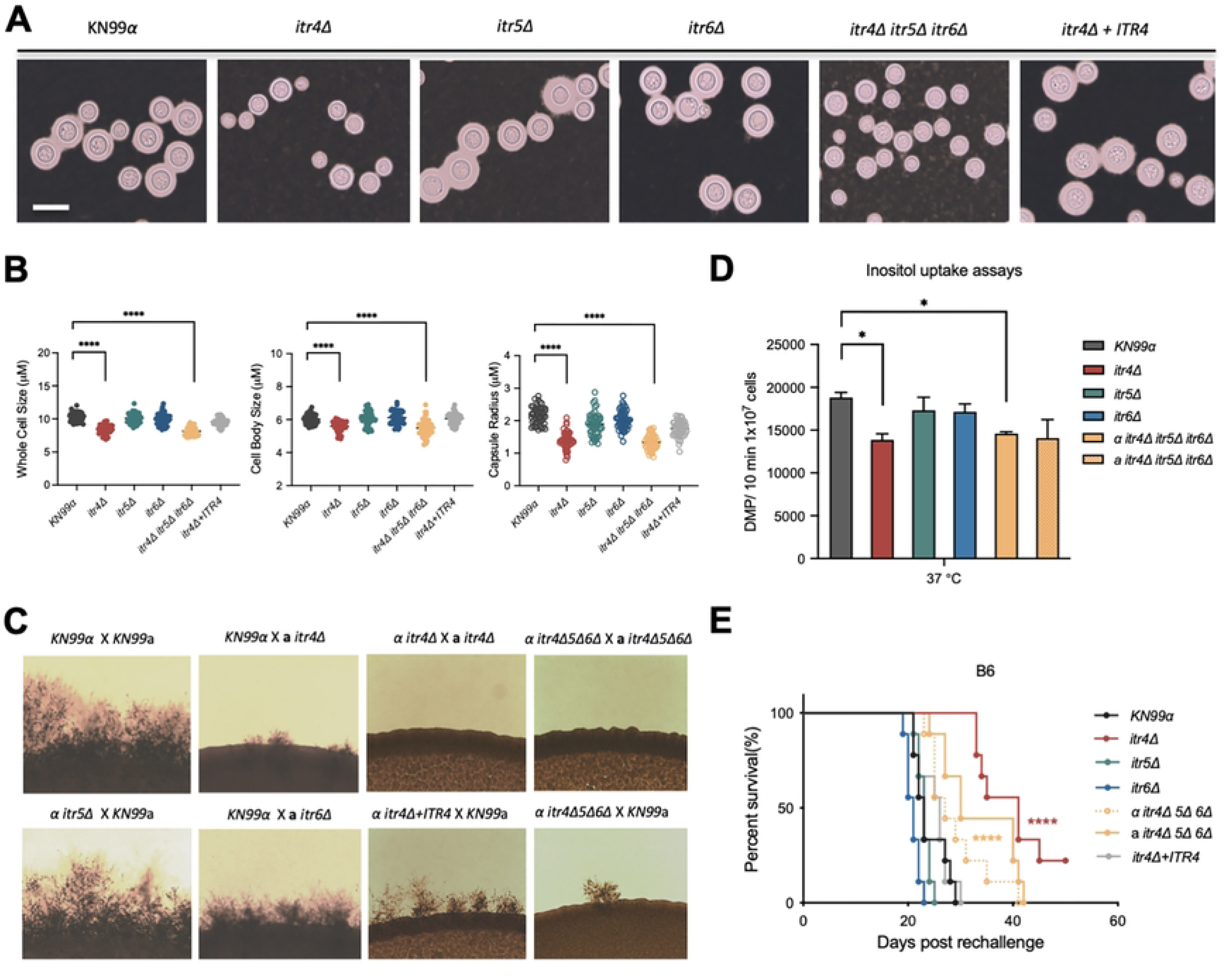
Analysis of group II *ITRs* through *in vitro* phenotype*, in vivo* virulence and inositol uptake. (A) Cell morphology of KN99, *itr4Δ, itr5Δ,* and *itr6Δ* single mutants and their triple mutants (*itr4Δ itr5Δ itr6Δ*) after grown on DME medium for 3 days. (B) Quantitative data of both cell size and capsule size from over 100 cells from each sample are presented. Statistical analysis was done using two-tailed Student’s t test. ****, P<, 0.0001. Bar, 10 µm. (C) Unilateral and bilateral mating assays were performed in MS medium. Mating cultures were incubated at room temperature in the dark for 5 days before mating hyphe were photographed. (D) Inositol uptake analysis of *C. neoformans* strains. Yeast cells were mixed with 3^H^-labeled inositol and incubated at 30°C for 10 min in triplicate (repeated three times with similar patterns). Error bars indicate standard deviations of results from the three replicates. (E) Survival rates of mice infected with KN99, *itr4Δ, itr5Δ,* and *itr6Δ* single mutants and their triple mutants *itr4Δ itr5Δ itr6* in a murine intranasal inhalation model. Groups of 10 mice were infected with 1x10^4^ yeast cells each strain. For the animal survival data: ****, P<, 0.0001. Statistical analysis was done using a log rank (Mantel-Cox) test.

We also evaluated the possibility that group II *ITR*s could be involved in stress responses. In our *in vitro* phenotypic assays, the *itr* null mutants showed normal growth under osmotic stress (1.5 M NaCl or 1 M Sorbitol) and oxidative stress (3.0 mM H_2_O_2_) (S1C Fig). We also examined the cell integrity of *itr* mutants by applying chemicals that target cell integrity, including sodium dodecyl sulfate (SDS), calcofluor white (CFW), and Congo red. SDS disrupts the plasma membrane and lyses cells with a defective membrane, while CFW specifically binds to chitin and Congo red binds to glucan and chitin in the cell wall to disrupt cell wall integrity. Our results showed that the *itr4Δ* mutant had a growth defect on SDS but not CFW or Congo red, indicating a defect in plasma membrane integrity (S1D Fig).

### Itr4 regulates sexual reproduction

We previously found that inositol is important for stimulating *Cryptococcus* mating to help the fungus complete its life cycle and several *ITR*s in Group I are required for fungal mating[11, 26]. To determine the potential involvement of group II *ITR*s in fungal mating, we tested their single and triple mutants in both unilateral (mutant x wild type) and bilateral (mutant x mutant) mating assays on MS medium. Significant reduction of mating hyphal production and defects in sporulation were observed in the unilateral mating assays of the *itr4Δ* single mutant and *itr4Δ itr5Δ itr6Δ* triple mutant, while the mating was completely blocked in their bilateral mating assays. No obvious mating defect was observed in unilateral and bilateral mating assays of *itr5Δ* and *itr6Δ* single mutants. The mating defect of the *itr4Δ* mutant could be fully complemented (Fig 1C). These results demonstrate that Itr4 is essential for inositol-induced fungal mating.

### Itr4 regulates inositol uptake activity

To understand whether the mating and capsule formation defect of the *itr* mutants were related to the defect in inositol uptake activity, we performed inositol uptake assays for parental wild type H99 and the group II *itr* single mutants, as well as the *itr4Δ itr5Δ itr6Δ* triple mutant. Consistent with our previous report[21, 23], our results showed that the *itr4Δ* mutant had significantly reduced inositol uptake activity when compared to the reference H99. Similar to the *itr4Δ* mutant, the *itr4Δ itrΔ5 itr6Δ* triple mutant strains also manifested reduced inositol uptake activity, while *itr5Δ* and *itr6Δ* single mutants showed wild type level of inositol uptake activity (Fig 1D). These results also indicate that Itr4 is the main *ITR* in group II with important biological function.

### Itr4, but not Itr5 or Itr6, is required for full fungal virulence

Our previous studies showed that Itr4 is required for full cryptococcal virulence, and the outcome of infection in animal models is mouse strain dependent, due to difference in host immune responses[23, 24, 27]. Here we evaluated the potential involvement of every group II *ITR* in fungal virulence in a murine infection model of systemic cryptococcosis. Ten C57BL/6 mice per group were inoculated with 1 x 10^4^ yeast cells per mouse of each strain intranasally, and their survival rates were determined. Our results showed that while mice infected by the *itr4Δ* mutant, or the *itr4Δ itr5Δ itr6Δ* triple mutant had a delayed mortality, the ones infected with the *itr5Δ* or *it6Δ* mutants showed a survival rate similar to the wild type H99. The virulence attenuation exhibited in the *itr4*Δ mutant was fully rescued by reintroducing the *ITR4* allele (*itr4Δ ITR4*) in the same murine model (Fig 1E). These results demonstrate that Itr4 was required for full virulence, while the *itr5Δ* and *itr6Δ* single mutants did not have an obvious impact on fungal virulence. This data is consistent with our previous report that Itr4 is required for full fungal virulence in C57BL/6 mice[23]. Our previous studies also demonstrated that the *ITR4* gene showed a significant polymorphism in clinical isolates from patients with cryptococcal meningitis[23]. Therefore, Itr4 is the only *ITR* in Group II *ITR*s that plays a significant role in regulating inositol uptake and full fungal virulence. Consequently, we focused on the functional analysis of Itr4 to better understand the inositol regulation of *Cryptococcus* pathogenesis.

### Itr4 is an inositol transceptor regulating inositol sensing

To better characterize Itr4 function, we expressed *ITR4* in a *Saccharomyces* mutant strain with deletions of both *ITR1* and *ITR2* (*Sc itr1 itr2*) that lacks inositol uptake activity[21]. Full-length cDNAs of *ITR4* was cloned into the yeast expression vector pTH74 under the control of the *ADH1* promoter to generate a *ITR4:GFP* fusion construct. The strains expressing *ITR4:GFP* localized most of their GFP fluorescence signals to the plasma membrane. The control strain expressing the empty vector had diffuse GFP fluorescence signal in cytosol (Fig 2A).

**Figure 2.**
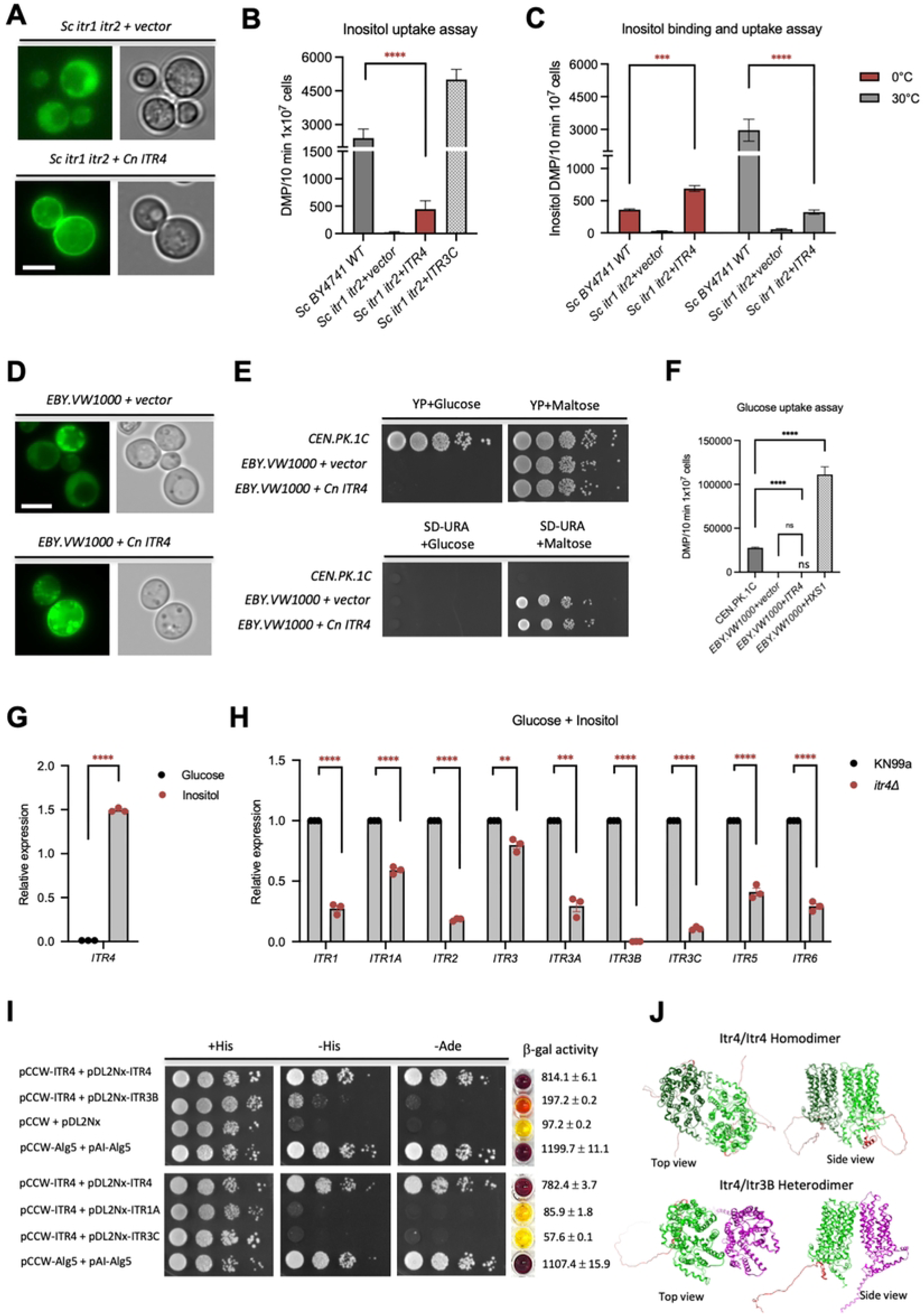
Heterologous expression and uptake assays of *ITR4* in *Saccharomyces* strains lacking inositol transporter. (A) Localization of *Itr4* in *S. cerevisiae* was determined by overexpressing a *Itr4:GFP* fusion protein in *Sc itr1 itr2*, a mutant lacking two inositol transporters. (B) Inositol uptake analysis of *Cryptococcus ITR* genes expressed in a yeast heterologous system was performed by detecting 3^H^-labeled inositol signal after 10 min incubation at 30°C. Error bars indicate standard deviations of data from at least three independent experiments. *Sc*, *S*. *cerevisiae*. (C) Inositol binding assay. Yeast cells were mixed with 3^H^-labeled inositol and incubated at 0°C for 10 min. (D) Localization of Itr4 in *S. cerevisiae* was determined by overexpressing a *Itr4:GFP* fusion protein in EBY.VW1000. EBY.VW1000 expressing an empty vector was used as a control. (E) Growth assay of *Saccharomyces* strains expressing *ITR4* on plates with indicated carbon sources. Cultures of the background strain CEN.PK2.1C and EBY.VW1000 expressing the pTH74 empty vector, or the *ITR4* gene were inoculated on SD media lacking Uracil but containing 2% maltose as carbon source. Serial 10-fold dilutions were prepared, and 5 µl of each dilution was spotted on different plates and incubated at 30°C or 37°C for 48 h before photography. (F) Glucose uptake assay was performed for CEN.PK2.1C and EBY.VW1000 expressing the empty vector, *HXS1* or *ITR4*. Yeast cells were mixed with 3^H^-labeled glucose and incubated at 30°C 10 mins. The error bar indicates the standard deviation of three repeats. (G) Quantification of the *ITR4* expression in a qRT-PCR assay. RNAs were prepared from KN99a cells cultured on medium with YNB+1% glucose or YNB+1% inositol for 4 hr. Values are expressed as relative expression of the *ITR4* gene, normalized to the *GAPDH* gene endogenous reference. (H) The expression of *ITRs* under KN99a and *itr4Δ* mutant strain background was determined in a qRT-PCR assay. (I) Detection of potential interactions between Itr4 and Itr1A, Itr3C and Itr3B in a yeast split-ubiquitin system. Two constructs, pCCW-Alg5 and pAI-Alg5, and pCCW and pDL2Nx serve as positive and negative control. Interaction was determined by the growth of yeast transformants on medium lacking histidine or adenine, and also by measuring β-galactosidase enzyme activities to verify the interaction. (J) Alphafold 2 protein modeling of Itr4-Itr4 homodimer and Itr4-Itr3B heterodimer structures.

We measured inositol uptake activity for the *Sc itr1 itr2* mutant strain expressing *C. neoformans ITR4* by using ^3^H-labeled *myo*-inositol (myo-[2-^3^H] inositol) at 30°C as described previously[21]. The yeast strain expressing *ITR4* had detectable signal but was significantly lower than the parental wild type BY4741 or the *Sc itr1 itr2* strain expressing the major inositol transporter *ITR3C* (Fig 2B), and close to the background noise. To determine whether the weak signal was due to inositol uptake or binding, we performed inositol binding assay at 0°C and detected a weak ^3^H signal in the *ITR4* yeast expressing strain that was similar to the signal detected in our uptake assay at 30°C (Fig 2C). This result showing a lack of inositol uptake in the heterologous system indicates that Itr4 may function as an inositol sensor (transceptor) rather than a typical transporter.

### Itr4 is not required for glucose uptake activity in a heterologous expression system

There was a possibility that Itr4 may transport other sugars, e.g., glucose. To examine whether Itr4 in *C. neoformans* functioned as glucose a transporter, the *ITR4:GFP* fusion construct was expressed under the control of the *ADH1* promoter in the *S. cerevisiae hxt* mutant strain EBY.VW1000, in which all 20 hexose transporters are deleted[28]. The EBY.VW1000 strain thus cannot grow on YPD medium but can grow on maltose as a carbon source. The expression of *ITR4* was confirmed by GFP signals and qRT-PCR. Our results showed that Itr4:GFP was localized on the plasma membrane as expected (Fig 2D).

The *S. cerevisiae* control strain CEN.PK2.1C, EBY.VW1000 expressing the empty vector or *P_ADH1_*-*ITR4:GFP* were used for growth assay on YPD or YP+Maltose. On YPD there was no growth of the EBY.VW1000 expressing empty vector or *ITR4:GFP*, while the growth defect was rescued on YP+Maltose condition, indicating the *ITR4* expressing strain can’t utilize glucose for their growth (Fig 2E). These data indicate Itr4 cannot take up glucose from medium to support yeast growth on YPD. To further confirm this, we examined the glucose uptake by Itr4 in this heterologous expression strain and found that no glucose uptake was detected in strains expressing *ITR4* or empty vector, while a control strain expressing *Cryptococcus* glucose transporter Hxs1 showed glucose uptake activity, confirming that Itr4 is not a glucose transporter (Fig 2F).

We posited that Itr4 may function as an inositol transceptor to regulate inositol uptake and metabolism. Such a sensory function might explain why the *itr4*Δ mutant, in which all other 9 *ITR*s remain present, showed significant defects in inositol uptake, virulence factors development, fungal mating, as well as fungal virulence (Fig 1). To test this hypothesis, we determined the potential role of Itr4 in regulation of other *ITR* family members by comparing the expression changes of remaining *ITR* genes in wild type H99 and its *itr4*Δ mutant. Our qRT-PCR results showed that the expression level of *ITR4* was highly induced in inositol condition compared to glucose condition (Fig 2G). Our qRT-PCR results also showed that most *ITR* genes had reduced expression in the *itr4*Δ mutant compared to wild type KN99, indicating that they were upregulated by Itr4 (Fig 2H). Together with its inositol binding activity shown in Figure 2C, our data identifies Itr4 as an inositol transceptor.

Receptors often form homodimer or heterodimer structure to sense signals and activate downstream pathways. To test the hypothesis that Itr4 may directly interact with other *ITR*s to form dimer structure to regulate inositol uptake, we performed protein-protein interaction assays using the split-ubiquitin system, a membrane-based yeast two-hybrid system to detect membrane proteins interactions[21]. We detected a strong interaction between Itr4 with itself, a weak interaction with Itr3B, and no interaction with two major inositol transporters Itr1A and Itr3C, based on both yeast growth on SD medium and ß-galactosidase activity (Fig 2I). These results indicate that Itr4 can form a homodimer and also a heterodimer with Itr3B, a gene that was silenced in the *itr4*Δ mutant background (Fig 2H). Putative protein dimer structure models were presented in Fig 2J.

In aggregate, our results demonstrate that Itr4 is an inositol-specific transceptor to sense inositol and regulate inositol uptake in *C. neoformans*. This assessment is consistent with the result that the *itr4*Δ single mutant showed significant defects in inositol uptake and exhibited strong inositol-related phenotypes.

### Itr4 regulates genes involved in inositol metabolism during inositol incubation

To further understand how Itr4 regulates *Cryptococcus* cells responding to inositol, we performed RNA-Seq assays to analyze transcriptional profiles of the *Cryptococcus* wild type KN99 and *itr4*Δ mutant that were either treated with or without inositol for 24 h. Our RNA-seq data identified a large number of differentially expressed genes (DEG) when compared between KN99 and the *itr4*Δ under YNB with either only 1% glucose or both 1% glucose and 1% inositol presence (Fig 3A). Compared to KN99, genes involved in inositol pyrophosphate biosynthesis and dephosphorylation, and inositol de novo biosynthesis were mostly upregulated in the *itr4*Δ mutant background, while genes involved in inositol uptake and inositol catabolism were down-regulated (Fig 3B), which is consistent with our qRT-PCR data (Fig 2H). *Cryptococcus* can use inositol as a carbon source. Conversion of inositol to glucuronic acid by inositol oxygenases (*MIOs*) is the first step of the only known pathway for inositol catabolism[25]. We found that the gene encoding inositol oxygenase 1 (*MIO1*) was induced significantly less by inositol treatment in *itr4*Δ than in KN99 (Fig 3B). These gene expression data demonstrate that Itr4 indeed positively regulates inositol acquisition and inositol catabolism.

**Figure 3.**
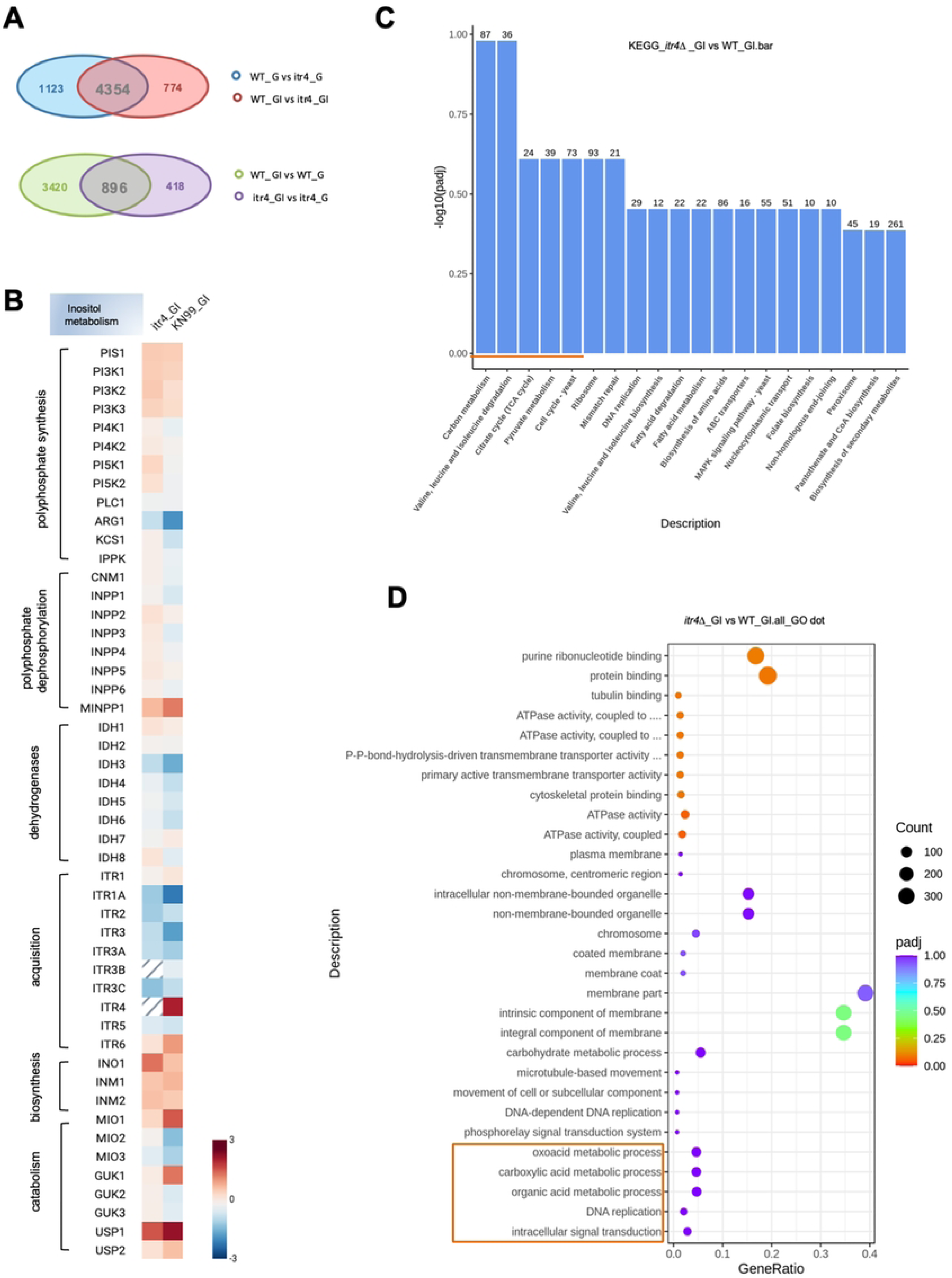
Functional Enrichment Analysis of Differentially Expressed Genes in the *itr4*Δ mutant. (A) Venn Diagram of Differentially Expressed Genes (DEGs) under different carbon conditions in the *itr4Δ* mutant vs. wild type. RNA-seq analysis identified DEGs (|log2 FC| > 0.5, p < 0.05) in the *itr4Δ* mutant versus wild-type KN99a strain under two conditions: YNB+1% glucose+1% inositol (glucose/inositol) and YNB+1% glucose (glucose-only). Venn diagram illustrating DEG distribution and overlap. Overlapping DEGs represent *ITR4*-dependent changes consistent across glucose-containing conditions, while unique DEGs in glucose/inositol reflect inositol-specific responses. (B) Expression profile of genes involved in inositol function. Intensity plot of genes involved in inositol synthesis, catabolism and metabolism. Gene expression of *itr4Δ* mutant and wild type KN99 were compared under condition of YNB+1%inositol+1%glucose as carbon sources. The scale bar ranges from +3 (red) to −3 (blue) Log2. (C) Bar plot of KEGG pathway enrichment analysis from RNA-seq data. Y-Axis (-log10(p.adj)) measuring statistical significance of pathway enrichment, adjusted for multiple testing (p.adj). Higher values indicate greater significance. X-Axis (Pathway Descriptions) listing enriched KEGG pathways, ordered from left (most significant) to right (least). Bar Length representing - log10(p.adj), longer bars denote higher significance. Numbers on Bars indicating the gene count and how many differentially expressed genes map to each pathway. Larger counts suggest broader involvement. (D) GO enrichment dot plot of top terms across Biological Process (BP), Molecular Function (MF), and Cellular Component (CC) categories. GO dot plot of selected terms, with dot size indicating gene count and color gradient showing significance. X-Axis (Gene Ratio) represents the proportion of differentially expressed genes in each term relative to the total genes annotated to that term. Higher ratios indicate a larger fraction of the pathway/term is impacted by the mutation. Dot Size (Count): Scales with the number of genes in the term. Larger dots mean more genes contribute to the enrichment, suggesting widespread effects. Dot Color (p.adjust) indicates adjusted p-value for statistical significance. Data from three biological replicates; enrichment performed using clusterProfiler (GO) and Pathview (KEGG) with H99 annotations from FungiDB.

Analyzing the differentially expressed genes (DEGs) using Ingenuity Pathway Analysis (IPA) identified several pathways that were upregulated by Itr4. A KEGG (Kyoto Encyclopedia of Genes and Genomes) pathway enrichment analysis highlighted the overrepresented pathways, including: ‘Carbon metabolism’ (map01200, p = 0.0018), ‘Valine, leucine and isoleucine degradation’ (map00280, p=0.0022), ‘Citrate cycle (TCA cycle)’ (map00020, p<0.0087), ‘Pyruvate metabolism’ (map00620, p = 0.0117), and ‘Cell cycle - yeast’ (map04111, p = 0.0155) (Fig 3C). A GO enrichment analysis plot (Fig 3D) revealed the top Biological Process (BP) terms include ‘DNA replication’ (GO:0006260, p < 0.01),‘intracellular signal transduction’ (GO:0035556, p < 0.05), and ‘organic acid metabolic process’ (GO:0006082, p < 0.01). Top molecular function (MF) terms included ‘protein kinase activity’ (GO:0004672, p < 0.05) and ‘sugar transmembrane transporter activity’ (GO:0051119, p < 0.05), that are linked to *PKC1* (CNAG_01845), *MPK1* (CNAG_04514), and *ITR3B* (CNAG_05667), identifying upregulation of proliferation, signaling, and metabolism as Itr4 regulated processes.

Interestingly, the glucose transceptor homolog gene *HXS1* (High affinity glucose transporter, CNAG_03772)[29] was significantly repressed (>125 fold) in the *itr4*Δ cells, while *SCW1* (cell wall integrity protein 1, CNAG_05663)[30], a gene required for cell wall integrity, and *BAT1* (Branched-chain-amino-acid transaminase, CNAG_05664)[30], a gene involved in amino acid metabolism, were completely silenced in the *itr4*Δ mutant background under the same culture condition (Table S4). These data may indicate a connection between inositol sensing and glucose uptake, cell wall integrity and amino acid metabolism, suggesting a cross talk among different metabolic pathways.

A volcano plot (Fig 4A) of the DEGs from the RNA-seq analysis of the *itr4Δ* mutant (itr4) versus the reference KN99 (WT) strain grown in YNB+1% glucose+1% inositol medium identified 5,128 DEGs, with 2,597 upregulated and 2,531 downregulated genes. Genes upregulated in the *itr4Δ* cells (red, right) include *PKC1*, *MPK1*, and *SNF1* (CNAG_06552)[31], genes associated with signal transduction, cell cycle, and metabolic pathways (Table S5), while downregulated genes (green, left) include *HXS1* and *MIG1* (CNAG_06327)[29, 32], reflecting disrupted glucose sensing and regulation (Table S4). Interestingly, a set of genes that exhibited near-zero expression in the *itr4*Δ (log2 FC<-5, p< 0.01) are all located near *ITR4* in the telomere region of chromosome 14, such as *SCW1*, *BAT1*, and *ITR3B*.

**Figure 4.**
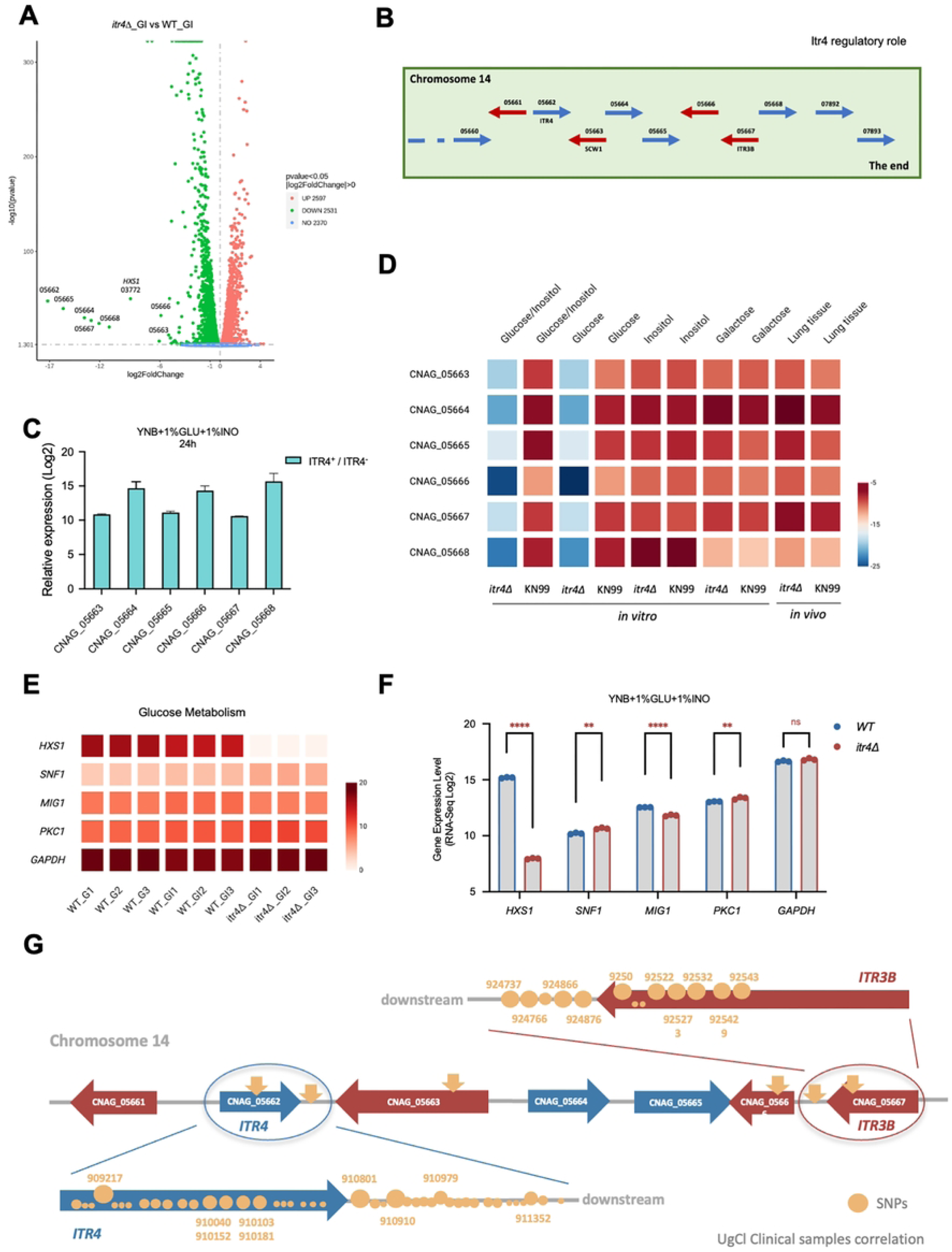
Gene silencing in chromosome 14 of *itr4*Δ induced by glucose. (A) Volcano Plot of DEGs from RNA-seq analysis of the *itr4Δ* mutant (itr4) versus wild-type KN99 (WT) strain grown in YNB+1% glucose+1% inositol. Volcano plot depicting log2 fold change (x-axis) versus -log10 adjusted p-value (y-axis, FDR-corrected). (B) Genomic organization of the *ITR4* gene locus on chromosome 14 in KN99a. The diagram illustrates a telomeric region containing *ITR4* and nearby genes. Blue arrows indicate rightward transcription (5’ to 3’ direction from left to right), while red arrows indicate leftward transcription (opposite orientation). ‘The end’ denotes the telomeric boundary. (C) qRT-PCR validation of transcriptional silencing in downstream genes on chromosome 14 in the *itr4Δ* mutant. The bar graph shows relative expression levels (normalized to wild-type) for genes located downstream of *ITR4* in glucose-containing conditions. Data represent mean ± SD from three biological replicates. (D) Heatmap of gene expression profiles of the chromosome 14 region under *in vitro* and *in vivo* conditions. Rows represent genes. Columns indicate conditions. Color scale denotes normalized expression levels nomolized with GAPDH (blue: low/silenced, -25; red: high, -5). Data from RNA-seq and qRT-PCR, averaged across replicates. (E) Heatmap showing normalized expression levels of *HXS1, SNF1, MIG1, PKC1,* with *GAPDH* as a reference, across wild-type (WT) cells grown in glucose alone (G1–G3), WT cells grown in glucose plus inositol (GI1–GI3), and *itr4Δ* cells grown in glucose plus inositol (GI1–GI3). (F) Quantitative data showing relative transcript abundance of CCR-associated genes comparing WT (1% glucose + 1% inositol) versus itr4Δ (1% glucose + 1% inositol). Data represent mean ± SD from three biological replicates. Statistical significance was determined using two-tailed Student’s t test, with asterisks indicating significance. (G) Schematic representation of single nucleotide polymorphisms (SNPs) in the *ITR* gene cluster on chromosome 14 from UgCl clinical isolates of *Cryptococcus neoformans*. The diagram represents the genomic organization of the telomeric region, with genes labeled by CNAG numbers: *ITR4* (CNAG_05662), *SCW1* (CNAG_05663), and *ITR3B* (CNAG_05667), along with flanking genes (CNAG_05661, CNAG_05664, CNAG_05665, CNAG_05666). Blue and red arrows indicate gene transcription orientation. Yellow arrows indicate SNPs identified in the *ITR* gene cluster on chromosome 14. Yellow circles represent percentage SNPs identified in clinical isolates, with numbers indicating their genomic positions (e.g., 909217). Upstream and downstream denote regions relative to the cluster. SNPs correlate with clinical outcomes, highlighting potential variants in *ITR4/ITR3B* associated with virulence or adaptation. Data from whole genome sequencing of UgCl cohort isolates.

### Glucose induced gene silencing in the *itr4*Δ background

In a recent GWAS study with *C. neoformans* clinical isolates, we found that *ITR4* and *ITR3B*, two *ITR* genes in the same telomere locus on chromosome 14, were significantly downregulated in three hypervirulent isolates that had single nucleotide polymorphisms (SNPs) in *ITR4*[23]. In a follow up study, several IFNγ-associated fungal genes (*PHS1*, *CNAG_05664, CNAG_06526*) had a greater than 2-fold change in expression in the *itr4*Δ strain compared to KN99[24]. Interestingly, based on our RNA-seq data, we found that all nine genes located outside of the *ITR4* in the telomere region of the chromosome 14, including (*CNAG_05663 (SCW1), CNAG_05664 (BAT1), CNAG_05665, CNAG_05666, CNAG_05667 (ITR3B), CNAG_05668, CNAG_07892* and *CNAG_07893*), were expressed in the wild type KN99 background, but not expressed in the *itr4*Δ mutant strain under high glucose condition (Fig 4B and 4C). Their expression was revived when the *itr4*Δ cells were cultured under medium lacking glucose but having other carbon sources (Fig 4D). These data indicate that glucose can induce a telomere silencing event in the cells lacking Itr4 but not in KN99 background. We also measured their expression during infection and found that these genes were all expressed in both the KN99 and *itr4Δ* background in the infected lung (Fig 4D).

### Itr4 regulates glucose metabolism through glucose sensor Hxs1

To better understand how *C. neoformans* was able to maintain the activity of both glucose metabolism and inositol metabolism under same high glucose conditions, we further examined the key CCR-associated genes in our RNA-seq data. In *S. cerevisae* glucose repression model, under high glucose condition, the inactivation of Snf1 kinase complex leads to activation of transcription repressor Mig1 in the nuclei to suppress alternative carbon utilization[33–36]. We found that in glucose only or glucose + inositol conditions, both *SNF1* and *MIG1* homologs were relatively stablely expressed, suggesting the glucose repression may not be as active under these conditions (Fig 4E). Striking, deletion of *ITR4* led to a significant repression of glucose sensor *HXS1* expression (>125 folds) relative to wild type under high glucose conditions, and genes involved in glucose uptake and metabolism are mostly downregulated, suggesting that Itr4 also regulates glucose sensing and transporting (Fig 4F and Table S6). It is possible that when *Cryptococcus* cell lacks inositol sensing due to *ITR4* deletion, the cell also slows down glucose utilization to maintain a balanced carbon utilization, leading to a glucose induced Itr4-dependent teleremore gene silencing as shown in Figure 4. Together, these data indicate that glucose does not simply suppress inositol-responsive transcription in wild type KN99, but instead requires Itr4 to maintain a balanced transcriptional state in which glucose and inositol metabolism are both active.

To investigate the clinical relevance of *ITR4* in cryptococcal infections, we analyzed whole genome sequencing data from a cohort of UgCl clinical isolates obtained from patients with cryptococcal meningitis in Uganda[24]. Interestingly, we found that multiple genes within the *ITR4*-associated chromosome 14 telomere region were also highly polymorphic, including *ITR4*, *SCW1*, CNAG_05666 and *ITR3B*. We identified abundant single nucleotide polymorphisms (SNPs) that were predominantly clustered around *ITR4* and *ITR3B*, with high levels of variation located downstream or within the *ITR4* ORF, and multiple variants located downstream or within the *ITR3B* ORF (Fig 4G). These variants were predicted to alter protein structure in Itr4, potentially affecting transmembrane domains, and suggesting possible regulatory impacts.

### Itr4 N-terminal tail is required for maintaining the Itr4 membrane localization and function

Given the potential for naturally occurring variants to influence the structure of Itr4, we evaluated the potential domains and amino acid sites required for Itr4 function. Utilizing the AlphaFold 2 structural modeling tool[37], we generated a model for the Itr4 structure (Fig 5A). Interestingly, protein structural modeling and protein sequence topological analysis showed that Itr4 has a unique long N-terminal cytosolic leader sequence (120 amino acids) before the first transmembrane domain (Fig 5A). Previous studies reported that glucose transceptors, e.g., Snf3 and Rgt2 in *S. cerevisiae*, have a long C-terminal tail that is required for their function as a glucose sensor by binding to membrane associated casein kinases[38]. To investigate the potential function of the Itr4 N-terminal leader sequence, we expressed an *ITR4* gene lacking this region (*ITR4^Δ11-119^*) in the *Sc itr1 itr2* mutant strain (Fig 5B). While the strain expressing *ITR4*:GFP showed plasma membrane localization, the strain expressing *ITR4^Δ11-119^* was no longer localized to the plasma membrane and instead showed an exclusively cytosolic GFP signal (Fig 5C). The inositol uptake assay also showed that the *Sc itr1 itr2* strain expressing *ITR4^Δ11-119^* did not have uptake activity (Fig 5D).

**Figure 5.**
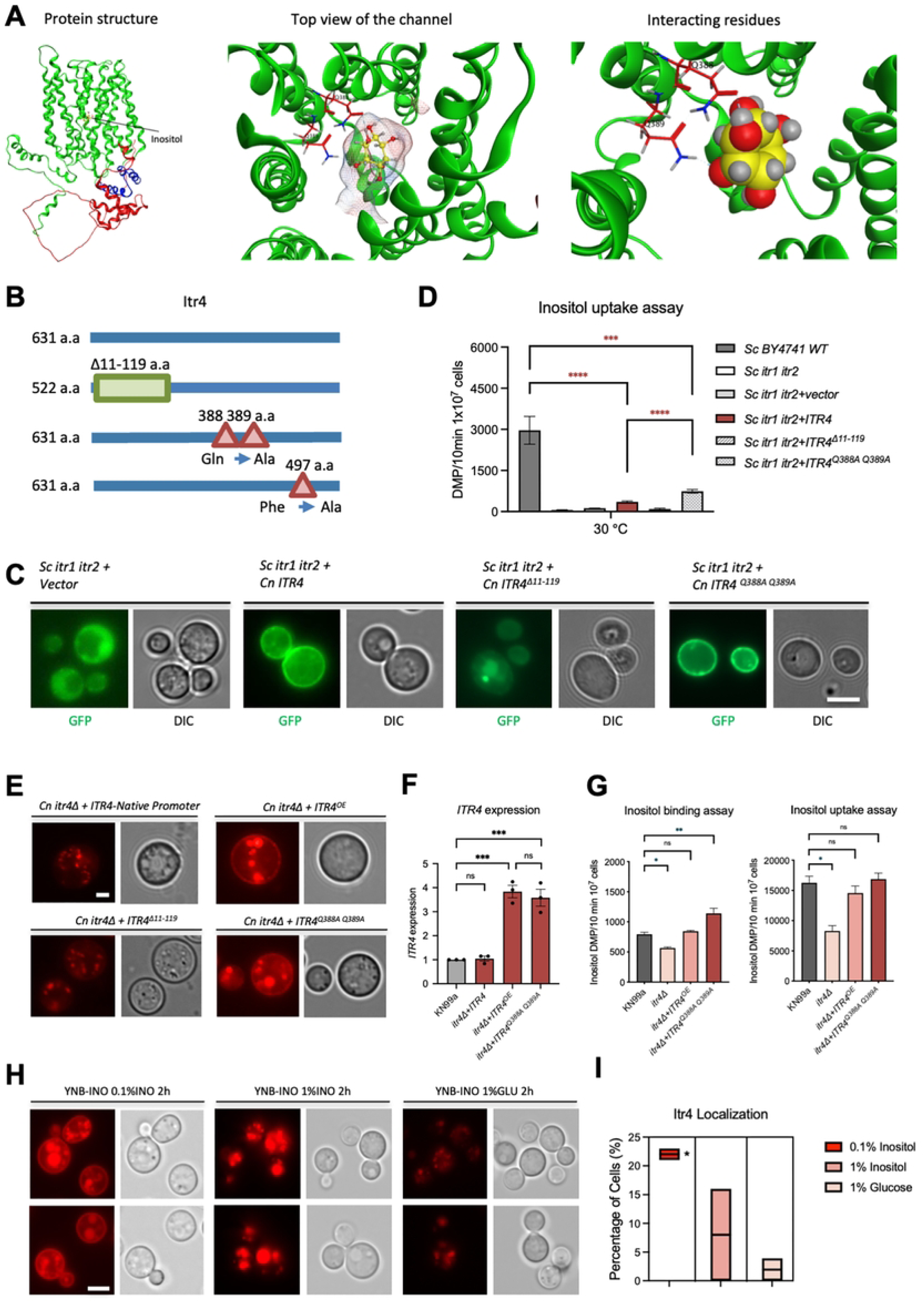
Itr4 protein structure modeling and predicted sugar binding sites. (A) Itr4 protein structure model predicted by AlphaFold2. Green color represents the Itr4 protein ribbon. Red region represents the first 120 amino acids of the N-terminual tail located within the cytoplasm. Blue region represents intracellular loops in Itr4. (B) Schematic of mutation sites in Itr4 protein. The N-terminal region between residue 11 and 119 amino acids was deleted, while residues Q388, Q389 and F497 were replaced by alanine (A). (C) Localization of *ITR4, ITR4^Δ11-119^, ITR4^Q388A, Q389A^,* and *ITR4^F497A^* in *S. cerevisiae* was determined by overexpressing of their GFP fusion protein in *Sc itr1 itr2* mutant lacking two inositol transporters. (D) Inositol uptake analysis of *Cryptococcus ITR4, ITR4^Δ11-119^* and *ITR4^Q388A, Q389A^* expressed in a yeast heterologous system. Yeast cells were mixed with 3^H^-labeled inositol and incubated at 30°C for 10 min. Error bars indicate standard deviations of data from at least three independent experiments. *Sc*, *S*. *cerevisiae*. (E) Localization of *ITR4, ITR4^Δ11-119^* and *ITR4^Q388A, Q389A^* in *C. neoformans* was determined by overexpressing of their mCherry fusion protein in *itr4Δ* mutant strain. (F) RT-PCR analysis of *ITR4* gene expression of different promoter. The *GAPDH* gene was used as a reference. (G) Inositol binding and uptake analysis of *Cryptococcus itr4Δ, ITR4^OE^,* and *ITR4^Q388A, Q389A^* strains. (H) Localization changes of *ITR4* in response of different carbon source. (I) Quantitative data of the percentage of cells with membrane localization.

In addition, expression of the *ITR4^Δ11-119^* in the *C. neoformans itr4Δ* mutant did not rescue the capsule defect phenotype, indicating a critical role for the N-terminal sequence in Itr4 function (Fig 6A). We posit that the N-terminal sequence is essential for sensing inositol and activating downstream signaling, a function similar to the C-terminal tail of Snf3 and Rgt2 in *S. cerevisiae*[38].

**Figure 6.**
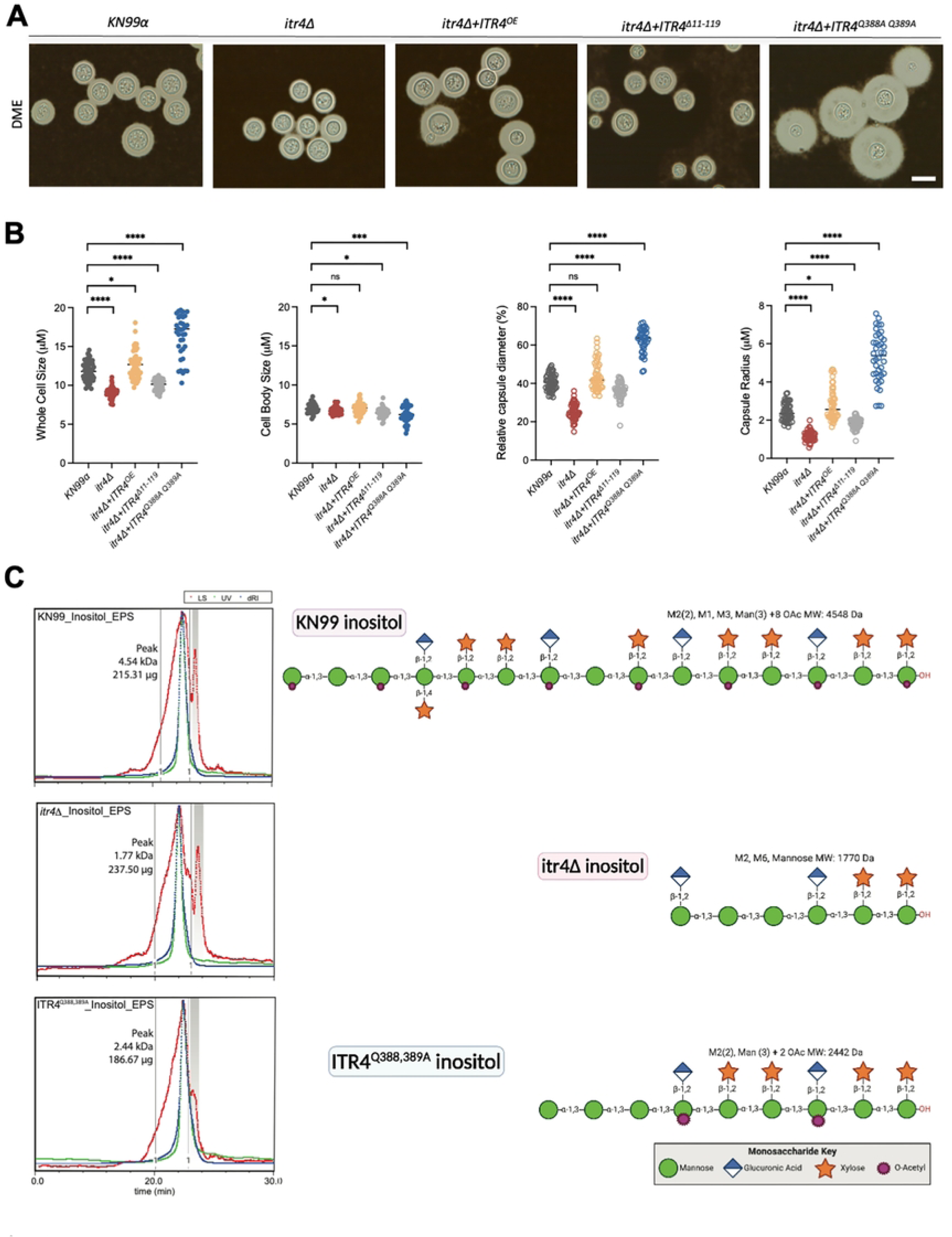
Itr4 regulates capsule size and structure. (A) Capsule formation of KN99, *itr4Δ, ITR4^OE^, ITR4^Δ11-119^* and *ITR4 ^Q388A,^* strains was visualized by India ink staining after cells were grown on DME medium for 3 days. (B) Quantitative data of both cell size and capsule size from over 100 cells from each sample are presented. Statistical analysis was done using two-tailed Student’s t test. ****, P<, 0.0001. Bar, 10 µm. (C) Characterization of GXM polymers in WT and the *itr4Δ* mutant strains. SEC-MALS analysis of EPS from KN99a, *itr4Δ,* and *ITR4 ^Q388A, Q389A^* cultured in inositol show differences in size. GXM motif composition of EPS given reported KN99a motif expression. These structures representative of but one of a number of ways these motifs could come together to make the polymer masses observed.

### Identification of key amino acids required for Itr4 function

Utilizing the Itr4 structural model, we further identified two predicted inositol binding sites (Q388 and Q389) (Fig 5A). To determine their potential importance for Itr4 function, we generated a *ITR4^Q388A, Q389A^* mutation allele and a *ITR4^F497A^* allele and expressed them in the *Sc itr1 itr2* heterologous system (Fig 5B). The F497 site was not directly interact with inositol but in close proximity in our structural model, and was selected as a control. Membrane localization was detected for both mutant alleles (Fig 5C). Inositol uptake assays showed that yeast strain expressing the *ITR4^Q388A, Q389A^* mutation allele had increased levels of inositol uptake activity compared to the strain expressing the wild type *ITR4* (Fig 5D), while the *ITR4^F497A^* allele did not exhibit any difference from while type allele (not shown). These data indicate that these two glutamine amino acids may function as a gate keeper to limit inositol transport, and their mutations partially open the channel. Our structure modeling also showed that the Q388A, Q389A mutation led to a large central cavity with lower interacting energy potential that may be more amenable to transport (S2 Fig). Therefore, the Q388A, Q389A point mutation produced a dominant active Itr4 allele that can both sense and transport inositol.

### The *ITR4^Q388A, Q389^*^A^ allele yields larger capsule but typical GXM particles

To further analyze the functional importance of Itr4 N-terminus and the putative inositol binding sites in *C. neoformans*, we generated mCherry tagged strains for Itr4, under the control of its native promoter (*P_ITR4_-ITR4:mCherry*). Utilizing the Actin promoter, we also generated strains overexpressing *ITR4* (*P_ACT1_-ITR4:mCherry*), *ITR4* lacking the N-terminus region (*P_ACT1_-ITR4^Δ11-119^:mCherry*) or *ITR4^Q388A, Q389A^* (*P_ACT1_-ITR4^Q388A, Q389A^:mCherry*) alleles. The expression and stability of *ITR4*, *ITR4^Δ11-119^* and *ITR4^Q388A, Q389A^* alleles were confirmed by the mCherry signal under fluorescence microscopy (Fig 5E). While the strains expressing *ITR4* or *ITR4^Q388A, Q389A^* retained membrane localization, the strain expressing the *ITR4^Δ11-119^* allele lost most of plasma membrane localization and instead most mCherry signal was in cytosol. The control strain expressing the *ITR4* under its native promoter manifested a weak plasma membrane fluorescence signal, suggesting a lower expression level compared to that under the control of the Actin promoter (Fig 5E and 5F).

We further measured inositol binding and uptake activities for the *itr4Δ* strain expressing of the *ITR4* and *ITR4^Q388A, Q389A^* alleles by using ^3^H-labeled *myo*-inositol (myo-[2-^3^H] inositol) at 0°C and 30°C as described previously[21]. Our results showed that the *itr4Δ* strain had lower binding and uptake signals than wild type strain KN99a, while the *itr4Δ* expressing *ITR4^Q388A, Q389A^* alleles had significantly higher binding activity than wild type strain (Fig 5G). This result indicates that the *ITR4^Q388A, Q389A^* allele may also enhance the inositol binding to the Itr4 ligand binding site in addition to increasing inositol transport activity, thus promoting inositol signaling activation.

To investigate the localization of fluorescence signal changes of Itr4 in response to different carbon sources, strains expressing *P_ITR4_*-*ITR4:mCherry* were grown in YPD overnight and reinoculated to YNB+0.1% inositol, YNB+1% inositol and YNB+1% glucose media and incubated for two hours. Our data clearly showed that, when *ITR4:mCherry* was under the Itr4 native promoter control, the majority of the mCherry fluorescence signal was weakly localized to the plasma membrane when cells grown on YNB+1% glucose, and its expression increased when switched to YNB+0.1% inositol medium. When cells were transferred to YNB+1% inositol condition, the fluorescent signal also increased, but was detected mostly in cytosol, indicating internalization of the Itr4:mCherry protein. The percentage of cells with membrane localization under different conditions was quantified (Fig 5H and 5I).

Consistent with our previous studies demonstrating that inositol can significantly affect the capsule[25], we found that the *itr4Δ* cells produced smaller capsule sizes than wild type cells (Fig 6A). To understand how Itr4 regulates capsule growth, we examined capsule in the *itr4*Δ mutant, the *ITR4^OE^* strain, the *ITR4^Δ11-119^* strain and the *ITR4^Q388A, Q389A^* strain under *in vitro* capsule inducing conditions[39]. We observed a significantly increased capsule size in the *ITR4^OE^* and *ITR4^Q388A, Q389A^* strains and a reduced capsule size in the *itr4*Δ and *ITR4^Δ11-119^* strain (Fig 6A and 6B). The large capsule size of the strain expressing the *ITR4^Q388A, Q389A^* allele is consistent with our hypothesis that it is a dominant active allele.

We further analyzed capsule structural changes in the *itr4*Δ mutant backgrounds by purifying the extracellular polysaccharides (EPS) secreted from the parent KN99a, *itr4*Δ and *ITR4^Q388A, Q389A^* strains. Our 1D [^1^H] NMR measurement showed no change in EPS structure or Glucuronoxylomannan (GXM) O-acetylation pattern between KN99a, *itr4*Δ and *ITR4^Q388A, Q389A^* strains. (S3 Fig). Further analysis by SEC-MALS (Size Exclusion Chromatography coupled with Multi-Angle Light Scattering) shows that while the structure of the EPS polymers did not change, the size of the polymers in both the *itr4Δ* and *ITR4^Q388A, Q389A^* strains were roughly half that of the parental strain (Fig 6C and Table S7). In addition, the *ITR4^Q388A, Q389A^* allele also showed less capsule density than either the parent or *itr4*Δ mutant. These data suggest that the EPS polymer size, rather than polymer structure, contributes to the changes in biological function of the *itr4*Δ mutant strains and their interactions with the host.

### Itr4 is required for *C. neoformans* to overcome the anti-fungal activity of macrophages

As a facultative intracellular pathogen, a key characteristic of *C. neoformans* is its ability to survive and replicate within the phagolysosome of macrophages[40]. The capsule has antiphagocytic effects during the macrophage-pathogen interaction[41]. To investigate whether Itr4 was required for fungal cell survival in macrophages, we performed *Cryptococcus*-macrophage interaction assays using the macrophage-like cell line J774. After coincubation with *C. neoformans* for 4 h, J774 cells were fixed with methanol and stained with Giemsa staining, and the phagocytosis rate was determined by the number of yeast cells inside each J774 cell. Macrophage-like cells co-cultured with the wild type KN99 resulted in ∼30% of J774 cells containing ingested yeasts, with ∼2.0 yeast cells on average per host cell. These phagocytosis numbers increased when the *itr4Δ* mutant or *ITR4^Δ11-119^* were co-incubated with J774 cells, but decreased when *ITR4^OE^* or *ITR4^Q388A, Q389A^* were co-incubated with macrophage-like cells, suggesting that Itr4 promotes phagocytic evasion (Fig 7A and 7B). To further test J774 killing of the fungal cells, *C. neoformans* and macrophage-like cells were co-incubated for another 4 or 24 h, and yeast survival rate was determined by colony forming units (CFUs). Our results showed that the *ITR4^OE^* and *ITR4^Q388A, Q389A^* strains had decreased responses to phagocytosis and were more resistant to J774 cell killing (S4A and S4B Fig). These results demonstrate that *ITR4* contributes to fungal survival in macrophages, likely by regulating the capsule size and structure. We found that the expression of the Th1 cytokine IFN-γ was significantly increased in the *itr4*Δ, *ITR4^OE^* and *ITR4^Δ11-119^* co-incubated macrophages (Fig 7C). These observations suggest that increased IFN-γ upon interaction is important during phagocytosis, which is consistent with our previously reported data[24].

**Figure 7.**
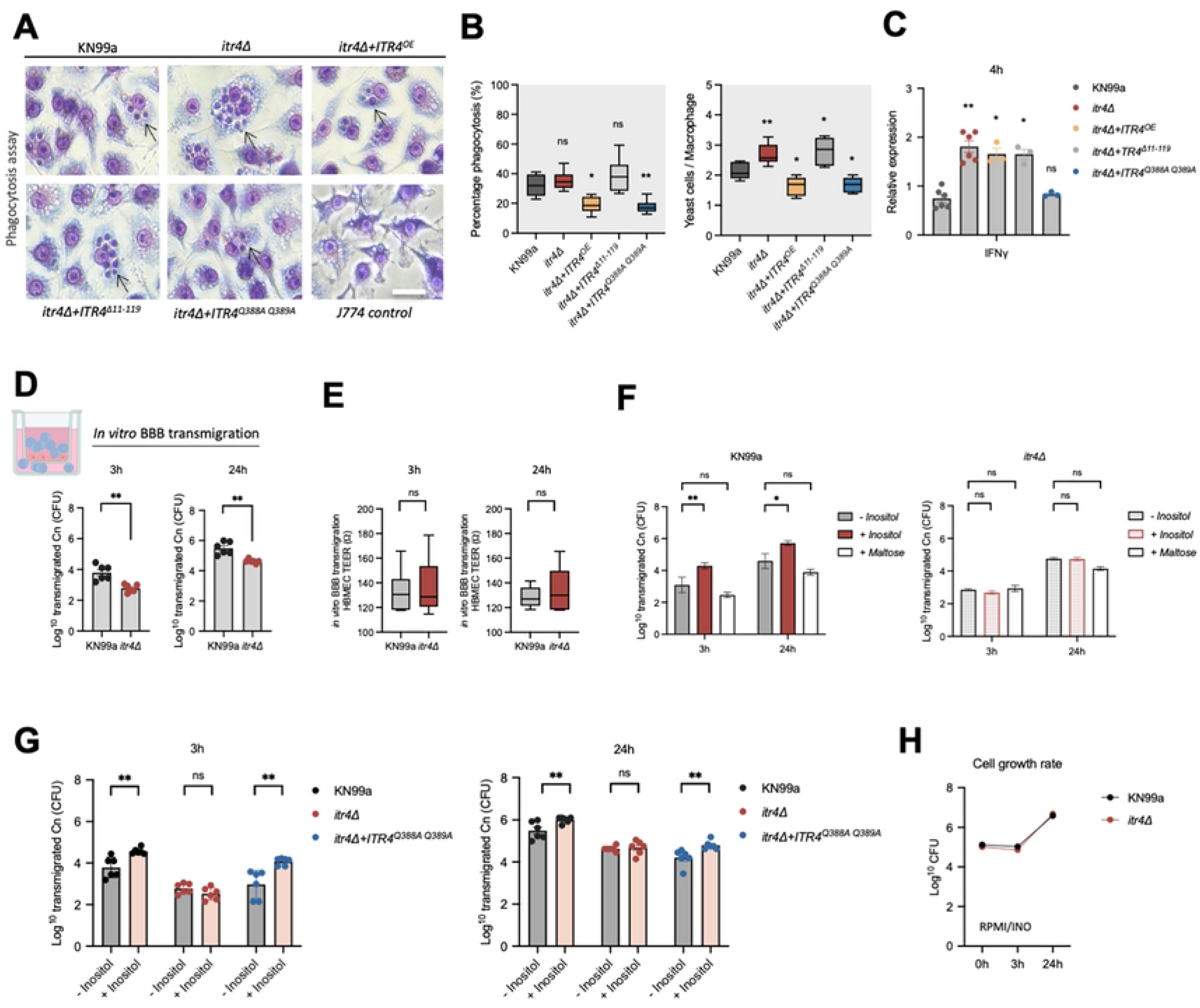
Itr4 is required for *C. neoformans* to overcome the antifungal activity of macrophages and cross the blood brain barrier. (A) Typical field views of phagocytosis assay for each strain. (B) The phagocytosis rate of *C. neoformans* in the J774 macrophage cell line was measured in 48-well plates. PBS-washed wild type KN99**a***, itr4Δ, ITR4^OE^, ITR4^Δ11-119^* and *ITR4^Q388A, Q389A^* strains were added to the activated macrophages and incubated for 4 h at 37°C with 5% CO2. Cells were fixed with methanol and stained with Giemsa stain before counting. The percentage of macrophages containing internalized yeast cells was measured from over 100 macrophages. Error bars indicate standard deviations. “**”, *P <* 0.01. “*”, *P <* 0.05 (determined by Mann-Whitney test). (C) Cytokine gene transcription during phagocytosis of each strain. Differential gene expression relative to GAPDH was examined by qRT-PCR with cytokine-specific primers and calculated by the ΔΔ*CT* method. The data shown are cumulative from three independent experiments. “**”, *P <* 0.01 (determined by Mann-Whitney test). (D) *In vitro* BBB transmigration assay. Cryptococcal transmigration assay was done with an *in vitro* human BBB model generated with HBMEC as described in Materials and Methods. Transmigration rate of wild type KN99 and *itr4*Δ mutant strains after 3 and 24 h incubation. (E) TEER resistance measurement of the Transwells with wild type KN99 and *itr4*Δ mutant strains. (F) Transmigration assays were done with *C. neoformans* strains KN99 and *itr4*Δ mutant strains in the absence or presence of 1 mM inositol in the bottom compartment. The equal concentration (1 mM) of inositol or maltose was added to the bottom compartment prior to addition of fungal cells to the top compartment. Maltose was used as a different monosaccharide control. The number of *Cryptococcus* cells transmigrated was determined by plating the medium collected from the bottom compartment after 3 and 24 h incubation. (G) Transmigration rate of indicated strains in the absence or presence of 1 mM inositol after 3 and 24 h incubation. (H) The grow rate of wild type KN99 and *itr4*Δ mutant strains in RPMI medium.

### Itr4 is required for inositol-mediated fungal traversal across the blood-brain barrier in an *in vitro* model

Previously we demonstrated that *C. neoformans* Itr1a and Itr3c are two major *ITR*s for inositol uptake and are required for dissemination across the BBB during brain infection[13]. To determine whether the inositol sensor Itr4 was required for fungal transversal across the BBB, we examined the transmigration ability of the *itr4Δ* mutant in the *in vitro* BBB model as previously described[13]. In the absence of additional inositol, transmigration of the *itr4Δ* mutant decreased by ∼50% at 3 and 24 h incubation period compared to the wild type, indicating that Itr4 is required for cryptococcal crossing of the BBB (Fig 7D). Under this condition, the Transendothelial Electrical Resistance (TEER) measurement indicated that the membrane was intact (Fig 7E). With inositol treatment, cryptococcal traversal was enhanced regardless of the presence of *ITR4*; however, the *itr4Δ* mutant showed significantly less transmigration than the parental control. The number of transmigrated wild type cells increased by ∼3-fold and ∼5-fold in the presence of inositol at 3 and 24 h incubation, respectively, compared to transmigration in the absence of supplemental inositol. In contrast, the *itr4Δ* mutant transmigration rate after 3 and 24 h was equivalent of the control (Fig 7F). We also tested the *ITR4^Q388A, Q389A^* strain for its transmigration efficiency using the *in vitro* BBB model. Our data showed that this mutant had reduced transmigration rate, likely due to its enlarged capsule size, but an increased transmigration rate was observed in the presence of additional inositol (Fig 7G).

Because inositol can be used as a carbon source for growth, there was a logistic concern that fungal cells with intact *ITR*s could grow better in medium with additional inositol. To address this concern, we compared the growth rate of wild type and the *itr4Δ* mutant in medium with or without addition of inositol. The results showed that all tested strains have similar growth rates regardless of the presence of additional inositol. (Fig 7H). Thus, the data are consistent with our interpretation that Itr4 plays an important role in responding to inositol availability and contributes to cryptococcal traversal across the BBB.

### The *ITR4^Q388A, Q389A^* allele shows attenuated virulence

*In vitro* analyses showed that Itr4 is important for capsule formation, mating, macrophage phagocytosis and BBB crossing, suggesting a potential role in host-pathogen interactions. We then examined the virulence of the wild type, the *itr4*Δ, *ITR4^OE^*, *ITR4^Δ11-119^* and *ITR4^Q388A, Q389A^* strains in a murine inhalation infection model. Our results showed that mice infected with *itr4*Δ, *ITR4^Δ11-119^* or *ITR4^OE^* all manifested prolonged survival than mice infected with wild type control, indicating a virulence attenuation. Interestingly, mice infected with the *ITR4^Q388A, Q389A^* strain had significantly delayed mortality, with a median survival of 45 d compared to 25 d with the reference strain (Fig 8A). These results indicate that the *ITR4^Q388A, Q389A^* allele had significant virulence attenuation.

**Figure 8.**
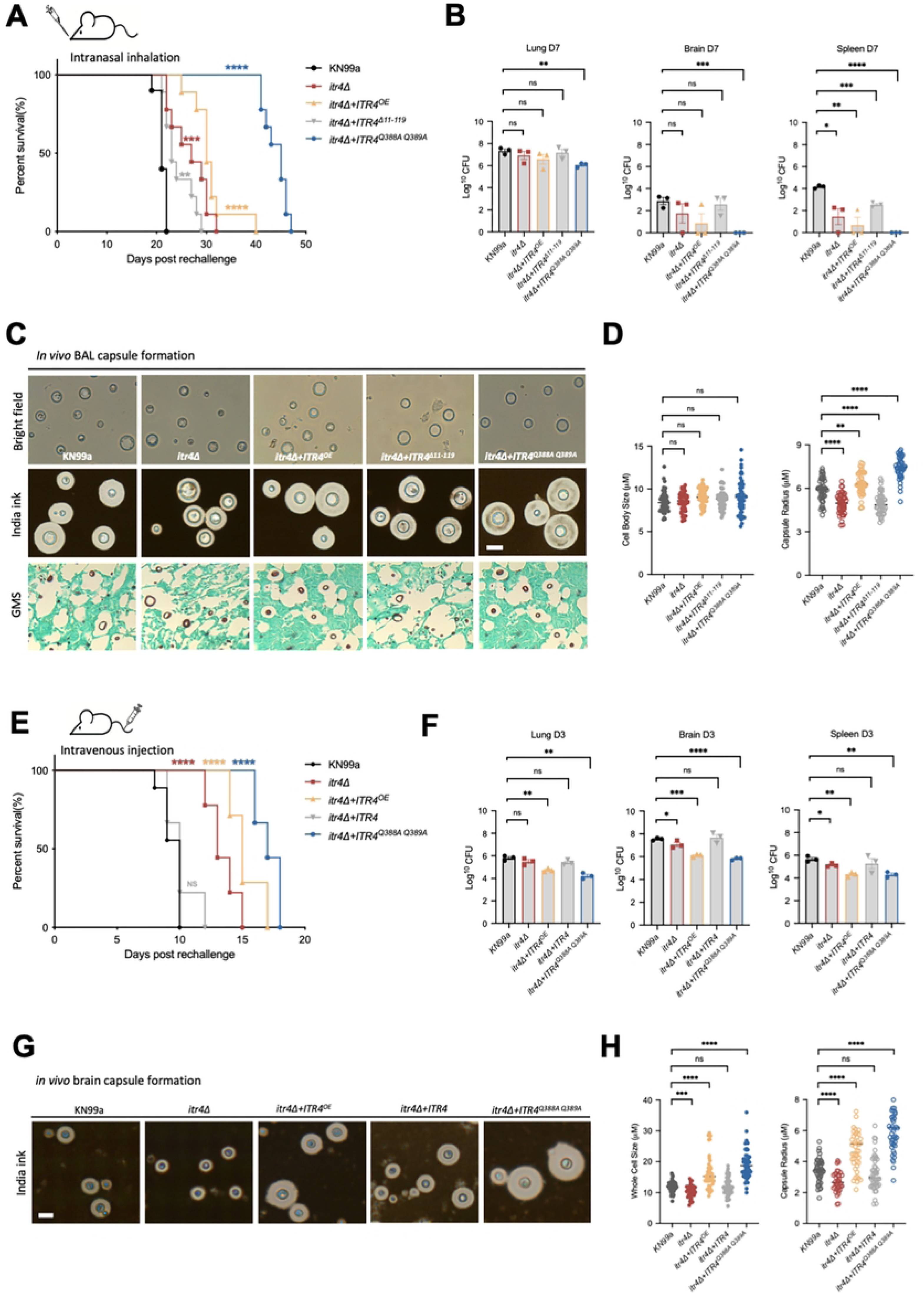
Itr4 is required for fungal virulence in animal models of systemic cryptococcosis. (A) Survival rates of wild type KN99 and its *itr4Δ, ITR4^OE^, ITR4^Δ11-119^* and *ITR4 ^Q388A, Q389A^* mutant strains in a murine intranasal inhalation model of systemic cryptococcosis. ****, P < 0.0001; statistical analysis was done using a log rank (Mantel-Cox) test. (B) Fungal burdens in the lungs, brains, spleens of animals infected with 1x10^5^ CFU of indicated strains at 7DPI. Each symbol represents one mouse. Bars represent the mean values +/- standard errors of the means. “****”, P *<* 0.0001; “*”, P *<* 0.001 “**”, P *<* 0.01; “*”, P *<* 0.05 (determined by One-way ANOVA nonparametric test for multiple comparisons). (C) BAL fluid and histopathological examination during lung infection at 7 DPI. These fungal cells were examined under microscope with or without India ink staining, and the white halo surrounding yeast cells reflects the capsule. For histological examination, the lungs were fixed and stained with GMS. Fungal cells appear dark brown or black with GMS staining. Yeast cells are highlighted (black arrow). (D) Quantitative results of yeast cell body size and capsule size during lung infection. For each of the five strains, 100 cells were quantified and conducted for statistical analysis based on the microscopic images. Each error bar indicates the standard deviation among the five independent quantifications. “**”, P < 0.01; “***”, P < 0.001 (determined by t test). (E) Survival rates of indicated strains in a murine intravenous injection infection model of systemic cryptococcosis. Balb/c mice were infected with 5x10^4^ yeast cells of each strain. “****”, P < 0.0001; statistical analysis was done using a log rank (Mantel-Cox) test. (F) Fungal burdens in the lungs, brains, spleens of animals infected with 5x10^4^ CFU of indicated strains at 3 DPI. (G) Brain tissue examination during intravenous injection infection. Cryptococcal cells were recovered from infected brain tissues at 3 DPI. These fungal cells were examined under microscope with India ink negative staining. (H) Quantitative results of yeast cell size and capsule size in the infection site in the brain in vivo. Quantitative data from over 100 cells from each sample are presented.

To better understand the mechanism of altered virulence of the *itr4*Δ mutant strains, three mice from each group were sacrificed at day 7 post-infection (7 DPI) and examined for fungal burdens in lungs, brains, and spleens. While significantly reduced CFU was recovered in the lungs infected with the *ITR4^Q388A, Q389A^* allele compared to wild type infected ones, no CFUs were recovered in brains and spleens in those animals, indicating a lack of dissemination at this timepoint (Fig 8B). Histopathology results of infected organs demonstrate less severe lesion development in lungs and brains infected by the *itr4Δ, ITR4^OE^* and *ITR4^Q388A, Q389A^* strains (S4 Fig). These observations are consistent with the virulence attenuation of these mutants. At the endpoint of the experiment, mice were examined for fungal burden in the lungs, brains and spleens. While all mice contained high levels of CFUs in the lungs, several of the *itr4Δ, ITR4^OE^*, *ITR4^Δ11-119^* and *ITR4^Q388A, Q389A^* strains were cleared of their brain or spleen infection, with some of them containing low levels of fungal CFUs compared to wild type control group (S4 Fig).

We isolated yeast cells from bronchoalveolar lavage fluid (BALF) of mice infected with different strains and analyzed their capsule size to examine the role of Itr4 in capsule size regulation during infection. Compared to wild type KN99 capsule in the BALF, we detected significantly increased capsule size in *ITR4^OE^* and *ITR4^Q388A, Q389A^* infected mice, which is consistent with the observations of *in vitro* conditions. Accordingly, the capsule size was smaller in the *itr4*Δ and *ITR4^Δ11-119^* cells during infection compared to the KN99a (Fig 8C and 8D). Altogether, these data confirm that Itr4 is involved in capsule size regulation during infection *in vivo*, and a significantly enlarged capsule may have slowed down the yeast dissemination, leading to virulence attenuation.

Because capsule is a major virulence factor that modulates host immune responses, we analyzed the expression of chemokine genes in lung tissues infected with different strains at 7 DPI by qRT-PCR. Uninfected C57BL/6 mice were analyzed as a control. We found that the expression of IFN-γ was significantly increased in the lungs of mice infected with *itr4*Δ, and significantly decreased in *ITR4^Q388A, Q389A^* infected mice (S4C Fig). These observations suggest that increased influx of IFN-γ upon infection with the *itr4*Δ mutant is associated with containment of the infection, which is consistent with our previously reported data[24]. Our lung gene expression analysis also showed that genotype-dependent differences in cytokine and macrophage polarization gene expression were coincided with fungal burden and survival outcomes. Compared to the parental strain, infection with the *itr4*Δ mutant strain showed higher expression of IFN-γ, IL-12a, IL-12b, TNF-α, and IL-17A, followed by increased *CCR2* and *NOS2* and lower *ARG1* expression. In contrast, the *ITR4^Q388A, Q389A^* strain showed reduced expression of these genes, with higher *ARG1* and IL-10 levels. IL-4 expression remained low across groups (S4 Fig and Table S8). Together, these data indicate that differential modulation of lung innate and chemokine responses distinguishes Itr4 alleles and aligns with their respective CFU and survival phenotypes in animal models.

### Involvement of *ITR4* in *in vivo* brain infection in a mouse intravenous model

Given that Itr4 was required for BBB crossing in our *in vitro* BBB model (Fig 7D and 7F), we evaluated the involvement of Itr4 in BBB crossing *in vivo*. Consequently, we infected mice intravenously with the *itr4*Δ, *ITR4^OE^*, and *ITR4^Q388A, Q389A^* strains. All these manifested significant virulence attenuation (Fig 8E), consistent with our observation in the inhalation animal infection model.

To determine the rate of fungal transversal across the BBB to cause brain infection in the intravenous infection model, mice injected with different *itr4*Δ mutant strains were sacrificed at 3 DPI; lungs, brains, spleens and kidneys were isolated and yeast CFUs were determined. Our results showed that there was a significant difference in fungal burden in the brain between mice infected by wild type and the *itr4*Δ mutant at 3 DPI. On the other hand, the fungal burden in all four organs was significantly decreased in *ITR4^OE^* and *ITR4^Q388A, Q389A^* infected mice compared to wild type (Fig 8F). Our results thus demonstrate that the inositol transceptor Itr4 is required for fungal cells to either cross the BBB or grow in the brain after transmigration.

To examine the role of Itr4 in capsule size control during CNS infection, we homogenized the brain tissues infected with different strains and analyzed their capsule size. Compared to wild type infected mice, we detected significantly enlarged capsule of the *ITR4^OE^* and *ITR4^Q388A, Q389A^* cells isolated from infected mouse brains, which is consistent with the observations of *in vitro* observations. Accordingly, the capsule size was smaller in the *itr4*Δ mutant during brain infection (Fig 8G and 8H). Histopathology of infected mouse brain tissue sections also indicated that the *ITR4^OE^* and *ITR4^Q388A, Q389A^* capsules are larger than the wild type and *itr4*Δ mutant strains (S4 Fig). Altogether, these data confirm that Itr4 is involved in capsule size regulation during infection *in vivo*, an observation consistent with in *in vitro* culture conditions. These results further confirmed that fungal Itr4 plays an important role in capsule size regulation and fungal pathogenesis, including *Cryptococcus* traversal of the BBB in CNS infection.

## Discussion

Pathogenic microbes must sequester nutrients from their hosts to support their proliferation and expression of virulence traits. We previously showed that *C. neoformans* can exploit the abundant inositol present in the brain for its development and virulence, supporting the progression of cryptococcal meningoencephalitis[12, 13, 21]. *C. neoformans* is an environmental fungus that occupies diverse ecological niches, such as plants and soils, where it interacts with amoebae. Since humans and animals serve as terminal hosts for *C. neoformans*, it has been proposed that the evolution of its virulence traits, and its capacity to infect human and animal hosts likely stems from adaptations acquired in its natural environments that also endow it with the capacity to resist clearance by the immune system. Previously we demonstrated that *Cryptococcus* has a highly sophisticated inositol utilization system, enabling it to efficiently acquire and metabolize inositol, which is abundant in plants such as *Eucalyptus* species, to complete its sexual cycle. This provides a potential explanation how this fungus produces infectious spores in nature[9, 11]. Interactions between *Cryptococcus* and soil amoebae (*Acanthamoeba castellanii*) have been shown to drive the development of virulence factors and confer the ability to survive within macrophages in mammalian hosts[42–44]. Similarly, the bacterial pathogen *Legionella pneumophila* can sequester inositol from amoebae, and its inositol metabolism promotes infection of both amoebae and macrophages[45]. It is conceivable that *C. neoformans* also senses and utilizes inositol in amoebae to support the development of fungal virulence factors, which helps survive predation. However, the mechanism by which *Cryptococcus* senses and responds to host inositol availability in the environment and in animal hosts was previously unknown.

In this study, we identified an inositol sensor, Itr4, which likely evolved from a transporter ancestor. Our data suggests that Itr4 enables the fungus to detect environmental inositol, thereby regulating the expression of genes involved in inositol acquisition and metabolism. This adaptive strategy, that presumably originally evolved for survival in the environment is also employed to efficiently sense and metabolize the abundant inositol found in the human brain. Therefore, we propose that this evolutionary specialization that once promoted survival in environmental hosts now facilitates infection of human hosts and that Itr4 serves as a critical sensor in this process (S5 Fig). To our knowledge, this is the first defined inositol sensor identified in eukaryotes. Whether other fungi possess a similar inositol sensor remains to be determined.

Although glucose is the preferred sugar and energy source for many pathogens, *C. neoformans* can efficiently take up and utilize free inositol, which is abundant in the brain, for its development and virulence, through its expanded ten member inositol transporter (ITRs) family and three member inositol oxygenases (MIOs)[13, 21, 25]. Transcriptomics studies have also shown that *ITRs* are upregulated in the cerebral spinal fluid (CSF) of patients with disseminated cryptococcal infections[46]. The identification of inositol sensor Itr4 in *C. neoformans* provides a mechanism on how the fungus senses and coordinates the function of this large family of ITRs. In this study, we further delineated the multifaceted role of Itr4 function, revealing its significance as a plasma membrane inositol transporter-like sensor (transceptor) and key regulator of inositol acquisition and metabolism. Our RT-qPCR and RNA-Seq data demonstrated that Itr4 regulates the expression of other *ITR*s, as well as genes involved in inositol metabolism (e.g., *MIO1*) in *C. neoformans*.

Sugar transceptors have been characterized in other fungi, including the well-studied glucose sensors Snf3 and Rgt2 in *S. cerevisiae*[35, 38]. These two sensors work in concert in the yeast CCR wherein when extracellular glucose levels are high Rgt2 is expressed and Snf3 is repressed, leading to expression of low affinity hexose transporter (e.g., Hxt1). Whereas in low glucose conditions Rgt2 is repressed and Snf3 is expressed inducing high affinity hexose transporters (e.g., Hxt2) expression[33]. Interestingly, while both Snf3 and Rgt2 have long C-terminal tails required for signal transduction, Itr4 contains a long N-terminal region essential for both membrane localization and function (Fig 5), with the *ITR4^Δ11-119^* allele failing to localize to the membrane and exhibiting the same capsule defect as that of *itr4*Δ (Fig 6). It is possible that this N-terminal region is responsible for interactions with downstream proteins, which remain to be identified. Based on structure modeling, we identified two amino acids required for Itr4 function. The *ITR4^Q388A, Q389A^* mutant allele showed increased inositol uptake activity, suggesting that those two glutamines prevent inositol transport into the cell by locking it in the channel. The Q-A Itr4 mutations result in functional inositol transport, leading to a dominant dual functional inositol sensor. The ammonium permease Mep2 is another transceptor with known dual function to sense and transport ammonium in yeast[47, 48]. Reported Mep2 protein structure revealed that the C-terminal region (CTR) is required for blocking ammonium transport[49]. Together these data indicate that Itr4 can sense but not transport inositol due to glutamine-medated channel restriction. It would be interesting to evaluate the function of similar amino acids in Snf3 and Rgt2 to see whether certain mutations could revive their glucose transporter function.

Our RNA-Seq data detected both glucose metabolism and inositol metabolism activities under a medium condition rich in both sugars (1% glucose and 1% inositol), suggesting regulation of sugar metabolism in *C. neoformans* is distinct from the classic carbon catabolte repression (CCR) model. It is established that yeasts like *S. cerevisiae* will utilize glucose before utilizing other carbon availability in the same medium through a glucose repression pathway that involves the glucose sensors Snf3 and Rgt2, kinase Snf1, and transcription repressor Mig1[34, 50]. However, our data suggests *C. neoformans* deviates from this model as fungal inositol metabolism remains active even in the presence of abundant glucose. The shift from glucose to inositol metabolism in *Cryptococcus* likely requires a nutrient sensor to determine the prevalence of inositol and activate transcription of the transporters. The cryptococcal hexose transcepter-like protein Hxs1 functions as a high-affinity glucose transporter with possible sensory function[29]. Our data shows that *ITR4* expression is induced while *HXS1* expression is downregulated in the presence of inositol even when abundant glucose is also present. Further, genes upregulated by inositol in the *itr4*Δ cells include metabolic enzymes (*IDH1*, *PDH1*) and DNA replication factors, while downregulated genes like *HXS1* are required for glucose transport and sensing, thus the presence of inositol alters metabolic pathways[51]. This metabolic flexibility, distinct from *S. cerevisiae*’s strict glucose repression, likely enhances *C. neoformans*’s host adaptation.

Beyond its regulation of inositol over glucose metabolism and the downstream effects of this, Itr4 also seems to regulate genes both within its transcriptional region and outside of it. Our RNA-seq and qRT-PCR show that *ITR4* deletion silenced genes outside of *ITR4* on chromosome 14 telomere region, including *ITR3B*, another inositol transporter, in a glucose-dependent manner. Interestingly compared to wild type, the *itr4*Δ mutant exhibited upregulation of *SNF1*, but downregulation of *HXS1* and *MIG1* indicating low glucose utilization mediated glucose derepression. Because Itr4 positively regulates inositol uptake and metabolism and also positively regulates glucose sensor Hxs1, our transcripome data suggest that the *itr4*Δ mutant had poor capacity to utlize these two carbons, which may lead to the glucose-dependent telomere silencing on Chromosome 14 in the *itr4*Δ mutant to limit metabolism and energy production, and to promote DNA replication and cell regeneration. This warrants further study on its silencing mechanism. Our findings position *ITR4* as a regulatory hub, sensing inositol to orchestrate glucose metabolism (*HXS1*), chromosome 14 telomere gene expression (*SCW1*, *ITR3B*), and downstream signaling (*PKC1*/*MPK1*, *SNF1*/*MIG1*). This dual role distinguishes it from other *ITRs* and mirrors nutrient-sensing transporters in pathogenic fungi[11]. The glucose-dependent silencing of downstream genes in the *itr4*Δ cells, which can be relieved by removing glucose, suggests *ITR4* may interact with a glucose-responsive regulator or chromatin modifier, a hypothesis that also warrants further exploration.

Through its receptor functions, we found that Itr4 is also involved in modulating the host response to *Cryptococcus*[25]. Our previous studies have demonstrated that inositol sensing and utilization influence capsule size and structure[24] which can, in turn, alter host responses like phagocytosis and pro-inflammatory IFNγ cytokine signaling. IFNγ plays an important role in host defense against *Cryptococcus* infection by regulating inflammation, activating macrophages and inducing T cell activation. In a recent genome wide association study (GWAS) of clinical isolates collected in Uganda[23, 24], we identified differences in inositol-related phenotypes*. ITR4* was one of several genes identified as having polymorphisms associated with differential human immune responses and patient survival. These results from clinical isolates further confirm the importance of the inositol sensor Itr4 in regulating fungal virulence and disease outcome in patients. Subsequent mouse model GWAS studies identified alleles in *ITR4* and a few other genes associated with increased production of IFNγ. We found that *itr4Δ* enhanced IFNγ production, while *ITR4* overexpression reduced IFNγ production.

Consistent with Itr4 playing a regulatory role in fungal virulence, the *ITR4^Q388A, Q389A^* mutation results in dramatically enlarged capsules while the *itr4Δ* capsule is significantly smaller than wild type. The *ITR4^Q388A, Q389A^* mutant also shows significantly decreased phagocytosis and elicited reduced IFNγ production while *itr4Δ* elicited increased IFNγ production. Further, in our *in vitro* transmigration assay and *in vivo* animal experiment, *itr4*Δ shows reduced BBB penetration and attenuated CNS dissemination, leading to delayed meningitis and prolonged survival. Surprisingly, both the *itr4Δ* and the *ITR4^Q388A, Q389A^* mutation result in attenuated virulence via intranasal and intravenous modes of infection. These discrepancies highlights Itr4’s role in balancing capsule size and functionality for infection and host survival suggesting that Itr4 plays a role in capsule regulation such that its mutation or loss results in dysregulation of capsule and therefore pathogenicity[25, 52]. These data led us to conclude that inositol sensing through Itr4 influences the capsule surface and subsequently both immune response and disease outcome.

In aggregate, Itr4 is a multifunctional transceptor critical for inositol sensing, cellular regulation, and pathogenicity in *C. neoformans*. Its loss triggers a cascade of molecular and phenotypic changes, revealing novel links between nutrient sensing, host adaptation, and fungal virulence (S5 Fig). Itr4 integrates glucose-inositol sensing to regulate capsule, phagocytosis, and virulence. While the mechanisms underlying glucose dependent chromosome 14 silencing in the *itr4*Δ mutant background warrants further investigation to fully elucidate the regulatory role of Itr4. The results expand our knowledge on the function of *C. neoformans ITR* family and highlight the unique contribution of Itr4 in nutrient sensing, host adaptation and fungal pathogenesis.

## Material and methods

### Strains, media, and growth conditions

*C. neoformans* strains used in this study are listed in Supplementary Table S1. *S. cerevisiae* strains and plasmids used in this study are listed in Supplementary Table S2. All primers used in this study are listed in Supplementary Table S3.

Strains were grown at 30°C on yeast extract-peptone-dextrose (YPD) agar medium and synthetic (SD) medium. YNB medium without inositol was purchased from MP Biomedicals. Niger-seed medium was used to test for melanin production. Dulbecco modified Eagles (DME) medium for assessing capsule production was prepared as previously described[9, 53]. A V8 juice agar medium (50 mL V8 original juice, 0.05% KH_2_PO_4_, and 2% agar, pH 5.0) and a modified Murashige and Skoog (MS) medium minus sucrose (Sigma-Aldrich) were used for mating and sporulation assays. Minimal medium (MM, 15 mM D-glucose, 10 mM MgSO_4_, 29.4 mM KH_2_PO_4_, 13mM Glycine, and 3.0 μM Thiamine. pH5.5) was used for capsule induction. All other media were prepared as described previously[26, 54, 55].

### Generation of *itr4Δ, itr5Δ* and *itr6Δ* mutants

Mutants for *ITR4, ITR5* and *ITR6* were generated in both H99α and KN99**a** strains backgrounds by overlap PCR as previously described[56]. The 5’ and 3’ regions of the *ITR4* gene were amplified from H99 genomic DNA with primer pairs CX1005 and CX1006, and CX1007 and CX1008, respectively. The 5’ and 3’ regions of the *ITR5* gene were amplified from H99 genomic DNA with primer pairs CX998 and CX999, and CX1000 and CX1001, respectively. The 5’ and 3’ regions of the *ITR6* gene were amplified from H99 genomic DNA with primer pairs CX991 and CX992, and CX993 and CX994, respectively (Table S3). Each target gene replacement cassette was generated by overlap PCR. Purified overlap PCR products were precipitated onto 10 µl gold microcarrier beads (0.6 µm; Bio-Rad), and strains H99 or KN99**a** were ballistically transformed as described previously. Stable transformants were selected on YPD medium containing G418 (200 mg/L) or NAT (100 mg/L). To screen for *itr4Δ*, *itr5Δ or itr6Δ* mutants, diagnostic PCR was performed by analyzing the 5’ junction of the disrupted mutant alleles with primers CX1009 and JH8994, CX1002 and JH8994, CX995 and JH8994. Positive transformants were identified by PCR screening with primers CX1010 and CX1011, CX1003 and CX1004, CX996 and CX997, respectively.

To generate complemented strains of the *itr4*Δ single mutant, a genomic DNA fragments that contained a 1.5 kb upstream region of *ITR4* ORF, the *ITR4* ORF, and its 500 bp downstream region was amplified by PCR with primers CX1222/CX1223. The *ITR4* PCR fragment was inserted into vector pJAF13 using in-fusion cloning. The construct *pJAF13-ITR4* was ballistically transformed in *itr4Δ* mutant strains. The selecting colonies grown on YPD with both nourseothricin and G418 were further confirmed by PCR. Phenotypic assays were performed to identify transformants in which the *itr4*Δ phenotype was complemented.

To generate overexpression strain of the *itr4Δ* mutant, the *ITR4* full-length genomic DNA was amplified with primers CX1800/CX1801 (Primer Table 3). The fragment was cloned into the BamHI site of vector pCXU200 containing the *Cryptococcus* actin promoter to generate a plasmid which contains *ITR4:mCherry* fusion fragment. Linearized plasmid was introduced into *itr4*Δ mutant strain to generate overexpression strain that expresses *Itr4:mCherry* protein.

### Generation of triple mutants and complemented strains

In mating assays, *C. neoformans* cells of opposite mating types were mixed and co-cultured on MS agar medium at 25°C in the dark for 10 days and filamentation was examined by light microscopy. Spore production was visualized by microscopy and photographed. Basidiospores were dissected from mating assays performed on MS medium.

To generate *itr4Δ itr6Δ* double mutants, a mating assay between α *itr4Δ::NAT* and **a** *itr6Δ::NEO* mutants was conducted and spores were isolated with an MSM system (Singer Instrument, England). Genomic DNA was isolated from all progenies that grew on YPD with both nourseothricin and G418. PCR was used to screen for *ITR4* gene deletion with primers CX1009/JH8994 and CX1010/CX1011, and for *ITR6* gene deletion with primers CX995/JH8994 and CX996/CX997 (Table S3). To generate *itr4Δ itr5Δ itr6Δ* triple mutants, a mating assay between α *itr4Δ::NAT itr6Δ::NEO* double mutant and **a** *itr5Δ::NEO* mutant was conducted and spores were isolated. Triple mutants were screened first by PCR on cultures initiated from single spores. All mutants were confirmed by PCR. The mating type of each confirmed mutant strain was determined by PCR using mating type specific primers and by genetic crossing.

### Assays for melanin and capsule production

Assays for melanin and capsule production were performed as previously described [53, 57]. In brief, melanin production was assayed by inoculating *C. neoformans* strains into 2 ml of YPD liquid medium, incubating the culture overnight at 30°C and spotting 5 µl of each culture with ten times series dilutions on Niger-seed agar medium. The agar plates were incubated at 30°C or 37°C for two days, and pigmentation of fungal colonies was assessed and photographed. To examine capsule production, 30 µl of overnight cultures were inoculated on DME agar medium and incubated at 37°C for three days. Capsule was visualized with India ink negative staining and observed with a 100x Olympus CX41 equipped with an Infinity digital camera (Olympus). Capsule sizes and cell body sizes of more than 100 cells were measured and calculated.

### Assays for stress responses and cell integrity

Each strain was incubated overnight at 30°C in YPD and sub-cultured in fresh YPD medium to OD_600_ ∼0.7. The cells were washed, resuspended, and serially diluted (1:10) in dH_2_O and spotted (5 μl) on YPD agar plates containing 1.5 M NaCl, or 1.0 M KCl for osmotic shock, or 2.5 mM and 5.0 mM H_2_O_2_ for oxidative stress. To test cell integrity, cells were also spotted on YPD agar plates containing 0.05% SDS, 0.5% Congo Red, or 20 μg/ml Calcofluor White (CFW). Plates were incubated at both 30°C and 37°C for two days and photographed.

### Inositol uptake assay

The inositol uptake assay was performed following a previously published method[19, 20]. In brief, the *Cryptococcus* strains were grown in YPD liquid cultures overnight at 30°C. Cells were diluted in YPD to an OD_600_ of 1.0, grown at 30°C, and collected at an OD_600_ of 5.0 by centrifugation at 2,600 *g* for 5 min. Cells were then washed twice with PBS at 4°C and resuspended in 2% glucose to reach a final concentration of 2 x 10^8^ cells/ml as determined by the use of a hemacytometer. For the uptake assay, the reaction mixture (200l) contained 2% glucose, 40 mM citric acid-KH_2_PO_4_ (pH 5.5), and 0.15 µM myo-[2-^3^H]-inositol (MP Biomedicals) (1 µCi/µl). An additional 200 µM concentration of unlabeled inositol (Sigma-Aldrich) was added to the reaction mixtures for competition assays. Equal volumes of the reaction and cell mixtures (60 µl each) were warmed to 30°C and mixed for the uptake assay, which was performed for 10 min at 30°C. As negative controls, mixtures were kept at 0°C (on ice) during the 10-min incubation. Aliquots of 100 µl were removed and transferred onto prewetted Metricel filters (1.2-µm pore size) on a vacuum manifold. The filters were washed four times each with 2 ml of ice-cold water. The washed filters were removed and added to liquid scintillation vials for measurements on a PerkinElmer TRI-CARB 2900TR scintillation counter.

### Glucose uptake assay

Full-length cDNAs of the *ITR4*, *ITR5* and *ITR6* genes were amplified from *C. neoformans* H99 total cDNA and were cloned into the yeast expression vector pTH74 to generate GFP fusion constructs, under the control of the *ADH1* promoter. *ITR4, ITR5* and *ITR6* expression plasmids were introduced into an *S. cerevisiae* strain EBY.VW1000 that lacks all 20 *HXT* transporters[28]. The expression of *Cryptococcus ITR4, ITR5* or *ITR6* in this yeast heterologous system was verified by both GFP localization and RT-PCR using gene-specific primers. Yeast strains were tested for growth on different medium at 30°C.

The *S. cerevisiae* control strain CEN.PK2.1C, the *S. cerevisiae hxtΔ* mutant strain EBY.VW1000, and EBY.VW1000 expressing empty vector, *ITR4:GFP*, *ITR5:GFP or ITR6:GFP* genes from *C. neoformans* were grown in YP containing 2% maltose liquid cultures overnight at 30°C. Collected cells were washed with dH_2_O, resuspended in YP containing 2% maltose, and incubated for 2 h. Then cells were suspended in PBS at a final concentration of 2 x 10^8^ cells/ml for uptake assay. Each 100 μl cell suspension was mixed with 100 μl labeled glucose (^3^H-glucose) solution at room temperature. Samples (100 μl) were removed after 30s, 1 min, 5 min, and 10 min, and mixed with 1 ml ice-cold water to stop the reactions. Cells were immediately collected on fiber filters, washed three times with 10 ml of ice-cold water, and transferred to scintillation vials for measurement.

### Detection of *ITR* gene expression using qRT-PCR

To test how the *ITRs* respond to the presence of environmental myo-inositol, both *in vitro* and *in vivo*, we measured the mRNA levels for *ITR* genes under different conditions via quantitative real-time PCR (qRT-PCR). Cultures of *C. neoformans* wild type strain H99 and its mutant strains were grown on YPD for 24 h at 30°C. Collected cells were washed with dH_2_O and pellets were used for total RNA extraction. Total RNAs were extracted using Trizol reagents (Invitrogen) and purified with Qiagen RNeasy cleanup kit (Qiagen) following the manufacturer’s instructions. Purified RNAs were quantified using a Nanodrop instrument (Thermo Scientific). First strand cDNAs were synthesized using a Superscript III cDNA synthesis kit (Invitrogen) following the instructions provided by the manufacturer. Expression levels of *ITRs* and *GAPDH* were analyzed with the comparative C_T_ method using Brilliant SYBR green QPCR reagents (Takara) as described previously [11, 21].

### Protein-protein interaction assay using the split-ubiquitin system

The split-ubiquitin system was utilized to investigate the interaction between Itr4 and Itr1A, Itr3C and Itr3B as previously described[54]. Vectors and yeast strains were included in the DUALmembrane Kit 2 (Dualsystem Biotech, Switzerland). *ITR4* full-length cDNA was cloned into yeast expression vector pCCW (the N-terminal half of the ubiquitin Cub protein was fused to the C-terminus of Itr4), and *ITR1A*, *ITR3C* and *ITR3B* full-length cDNA was cloned into the pDSL-XN vector (the mutated C-terminal half of ubiquitin NubG protein was fused to the *ITR1A*, *ITR3C*, *ITR3B* and *ITR4* N-terminus). All cDNA sequences were confirmed by DNA sequencing. Cub and NubG fusion constructs were co-transformed into host yeast strain NMY32. Two constructs, pCCW-Alg5 and pAI-Alg5, and pCCW and pDL2Nx serve as positive and negative control. Interaction was determined by the growth of yeast transformants on medium lacking histidine or adenine, and also by measuring β-galactosidase enzyme activity assays using Chlorophenolred-β–D-galactopyranoside (CRPG; Calbiochem, San Diego, CA) as substrate.

### Fluorescent signal quantification and data analysis

Plasmids expressing *P_ACT1_-ITR4:mCherry* fusion protein, *P_ACT1_-ITR4^Δ11-119^:mCherry* fusion protein and *P_ACT1_-ITR4^Q388A,Q389A^:mCherry* fusion protein were linearized and transformed into *itr4*Δ mutant to generate the *itr4*Δ *ITR4^OE^, itr4*Δ *ITR4^Δ11-119^* and *itr4Δ ITR4^Q388A, Q389A^* strains. Strains were cultured in YPD overnight and were observed with a Nikon fluorescence microscope.

The percentage of mCherry signals localized within the cell (% cyto-sollocalized GmCherry) was calculated in accordance with data analysis seen in Hankins *et al*[58]. The outside of each cell was outlined using the freehand selection tool in ImageJ and then the total fluorescence of the cell was measured (Fluor_total_). The area just within the plasma membrane was then outlined to give the total internal fluorescence excluding the plasma membrane (Fluor_int_). The percentage of fluorescence localized within the cell was found by dividing Fluor_int_ by Fluor_total_.

### *Cryptococcus*-macrophage interaction assay

The *Cryptococcus-*macrophage interaction assay was done as previously described[59]. Macrophage-like J774 cells were cultured in DME medium with 10% heat-inactivated FBS at 37°C with 5% CO_2_. J774 cells (5x10^4^) in 0.5 ml fresh DME medium were added into each well of a 48-well culture plate and incubated at 37°C in 5% CO_2_ overnight. To activate macrophage cells, 50 units/ml gamma interferon (IFN-γ; Invitrogen) and 1 µg/ml lipopolysaccharide (LPS; Sigma) were added into each well. *C. neoformans* overnight cultures were washed with phosphate-buffered saline (PBS) twice and opsonized with 20% mouse complement. *Cryptococcus* cells (2x 10^5^) were added into each well (yeast/J774 ratio, 4:1). To assess the phagocytosis rate, the cells were washed with PBS after a 1.5 h coincubation and fixed with methanol for 30 min. Giemsa stain was added to the wells at a 1:5 dilution, and the plates were incubated overnight at 4°C. Cells were washed once with PBS and analyzed using an inverted microscope. To assess intracellular proliferation of *C. neoformans*, nonadherent extracellular yeast cells were removed by washing with fresh DME medium after a 4 h coincubation and cultures were incubated for another 0, 4, and 24 h. At indicated time points, the medium in each well was replaced with distilled water (dH_2_O) to lyse macrophage cells for 30 min at room temperature. The lysate was spread on YPD plates, and CFU were counted to determine intracellular proliferation.

### Human brain microvascular endothelial cells (HBMEC) transmigration assay

Primary isolates of HBMEC were cultured as previously described[60]. HBMEC were routinely grown in RPMI 1640 supplemented with 10% heat-inactivated fetal bovine serum, 10% Nu-serum, 2 mM glutamine, 1 mM sodium pyruvate, penicillin (100 units/ml), streptomycin (100 mg/ml), essential amino acids, and vitamins. The cells were incubated at 37°C in a humidified incubator with 5% CO_2_. Before each experiment, the culture medium was replaced with experimental medium containing Hams-F12/M199 (1:1, v/v), supplemented with 5% heat-inactivated fetal bovine serum.

The *in vitro* human BBB model was generated and used for fungal transmigration assays as previously described[61, 62]. HBMEC were seeded on Transwell polycarbonate tissue culture inserts with a pore diameter of 8.0 μm (Corning Costar) and cultured until their trans endothelial electrical resistance (TEER) reached over 350 V/cm^2^, as measured by an Endohm volt/Ω meter in conjunction with an Endohm chamber (World Precision Instruments). The medium was replaced with experiment medium before each experiment. Yeast cells were washed with phosphate-buffered saline (PBS) and resuspended in HBMEC culture medium. 10^5^ Cryptococcus cells were added to the top compartment and then incubated at 37°C. At 3 and 24 h, the medium in bottom compartments was collected and immediately replaced with fresh medium. Fungal cell numbers in the collected medium were addressed by CFU counts to determine the number of transmigrated viable yeast cells. To determine the specificity of myo-inositol, the transmigration assay was performed in the presence of myo-inositol, D (+) galactose or D (+) maltose (1 mM each) in the bottom compartments prior to addition of Cryptococcus cells (10^5^) in the top compartment of Transwells. Results are presented as the total number in the bottom chamber. Each set was triplicated and repeated three times independently[13]. The statistical analysis of the data from our in *vitro* studies was done with a two-tailed Student *t* test. Statistical significance was determined at P, <0.001.

### RNA-seq Quantification Analysis

*C. neoformans* cells of the WT and *itr4Δ* mutant strains were cultured overnight at 30°C in YPD liquid medium and then inoculated in YNB+1% Glucose+1% Inositol condition for 24 h. The cells were then collected and total RNAs were extracted using Trizol reagents (Invitrogen) and purified with MACHEREY-NAGEL RNA purification kit following the manufacturer’s instructions. Purified RNAs were quantified using a Nanodrop instrument (Thermo Scientific). RNA-Seq analysis was performed by Novogene according to the company’s protocol (https://www.novogene.com/us-en). For the RNA-seq, cDNA library preparation, Illumina sequencing, and data analysis were performed by Novogene (Sacramento, CA). Sequencing was performed using Illumina NovoSeq 6000 to generate 150 bp paired-end reads. Approximately 50 million raw reads were generated. Adapter and low-quality reads or reads with uncertain nucleotides constitute more than 10 percent of either read (N>10%), and reads with low quality nucleotides (base quality less than 5) constitute more than 50 percent of the read, were removed. Clean reads were mapped to the annotated genome of *C. neoformans* H99 (NCBI:txid235443) using HISAT2 software (version 2.0.5). Differential expression analyses were conducted using the DESeq2 package (version 1.20.0)[63]. Significance was determined using multiple unpaired T-tests with two stage linear step-up.

## Data Availability

All raw RNA-seq data were deposited at NCBI Sequence Read Archive (SRA) (https://www.ncbi.nlm.nih.gov/sra) with BioProject ID PRJNA1142147[https://www.ncbi.nlm.nih.gov/sra/PRJNA1142147]. Processed signature data can be accessed in supplementary tables (Table S4 – S6). The STRING (Search Tool for Retrieval of Interacting Genes/Proteins) database can be found at http://string-db.org. Source data are provided with this paper.

## Murine infection

Survival curves of infected mice in a *Cryptococcus* murine inhalation model as previously described[21, 25]. *Cryptococcus* strains were grown at 30°C overnight and cultures were washed twice with 1x phosphate-buffered saline (PBS) by centrifugation and resuspended at a final concentration of 2 x 10^5^ CFU/ml. Groups of 10 female C57BL/6 mice (Jackson laboratory, ME) were used for each infection. For the intranasal inhalation model, mice were intranasally infected with 10^4^ yeast cells of each strain in 50 µl PBS as previously described. For the intravenous injection model, 5 x10^4^ yeast cells in 100µl volume for each strain were inoculated via tail vein injection. Over the course of the experiments, animals that appeared moribund or in pain were sacrificed by CO_2_ inhalation. Survival data from the murine experiments were statistically analyzed between paired groups using the long-rank test and the PRISM program 6.0 (GraphPad Software) (P values of <0.001 were considered significant).

## Isolation and measurement of yeast cells from mouse BALF

Yeast strains were grown at 30°C overnight and cultures were washed twice with PBS buffer and resuspended to a final concentration of 2×10^7^ cells/mL. Groups of 5 female C57BL/6 mice were intranasally infected with 1×10^6^ yeast cells of each strain as previously described[64, 65]. BALF samples were harvested at day 3 after inoculation. BALF was collected in 3 ml of 1× PBS buffer using a catheter inserted into the trachea of animal post-euthanasia, and airway-infiltrating cells were lavaged with ∼0.7 ml of 1× PBS at one time to a total volume of 3 ml. Cells were spun down and host cells were lysed by adding sterile ddH_2_O and incubating for 45 min at room temperature. *Cryptococcal* cells were examined under microscope. Capsule sizes and cell body sizes of more than 500 cells were measured and calculated.

## Histopathology and immune response analysis for infected organs

Infected animals were sacrificed at the end of the experiment according to the Rutgers University IACUC approved animal protocol. To compare the fungal burdens and host inflammatory responses, the lungs, brains, and spleen were dissected and fixed in 10% formalin solution for section preparation at Rutgers University Histology Core Facility. Tissue slides were treated either with hematoxylin and eosin (H&E) staining for bronchus-associated lymphoid tissue, or with Grocott’s methenamine silver (GMS) staining for fungal morphology observation *in vivo*. Infected lungs, brains, and spleens were also isolated and homogenized (Ultra-Trra T8, IKA) in 3 ml cold 1 x PBS buffer for 1 minute for each type of organ. The tissue suspensions were serially diluted and plated onto YPD agar medium with ampicillin and chloramphenicol, and colonies were counted after 3 days of incubation at 30°C.

Lung tissue total RNAs were extracted using Trizol reagents (Invitrogen) and purified with a RNeasy cleanup kit (Qiagen) following the manufacturer’s instructions. Purified RNAs were quantified using a Nanodrop instrument (Thermos Scientific). First strand cDNAs were synthesized using a Superscript III cDNA synthesis kit (Invitrogen) following the instructions provided by the manufacturer. Expression levels of Cytokines and *ACTIN* were analyzed with the comparative C_T_ method using Brilliant SYBR green QPCR reagents (Takara) as described previously Relative mRNA levels were determined by quantitative reverse transcription qRT-PCR. Gene expression relative to that of a naive sample was calculated by the ΔΔ*CT* method.

## In-Silico protein structure analysis

Structural modeling and ligand docking: The three-dimensional structures of Itr4 monomer and several dimers were predicted using AlphaFold2 in monomer and multimer modes respectively[37]. AlphaFold2 was accessed through the COSMIC2 portal. The resulting PDB files were transferred to the Rutgers University Amarel high-performance compute cluster for analysis. The structures were assessed in Molecular Operating Environment (MOE) version 2024.06 and the structures with the fewest clashing residues were prepared in MOE. The Amber:EHT forcefield was used for all operations within MOE. For docking, the Itr4 monomer and the energy-minimized myo-inositol were analyzed within MOE. The residues within the transmembrane helices were saved as sets and labeled TM1-12 according to their location. A dummy atom was created and residues within 4.5 Å were selected and the ligand-receptor pair with the lowest energy was selected. The resulting pose was captured as an instance within a MOE storyboard, imaged, and the ligand interaction diagram was reported. This storyboard instance was the starting point for subsequent virtual mutations.

Digital Alanine scan: The protein builder tool was used within MOE to mutate Q388 and Q389 to alanine as monosubstituted and disubstituted models. The protein builder minimization tool was used to minimize the energy of the selected residues that were substituted. A molecular surface representing the local electrostatic environment was produced using receptor atoms within 4.5Å of ligand atoms. The resulting mutant structure was saved as a new storyboard instance and the primary MOE viewer was reset to the original storyboard instance for a subsequent mutation.

## Ethics statement

The studies performed were governed by protocol 999901066 as approved by the IACUC committee of Rutgers New Jersey Medical School. Animal studies were compliant with all applicable provisions established by the Animal Welfare Act and the Public Health Services (PHS) Policy on the Humane Care and Use of Laboratory Animals.

## Acknowledgements

We thank David Meya at Makerere University for providing the Uganda clinical isolates. We thank Joe Heitman at Duke University for providing *Saccharomyces* and *Cryptococcus* strains. We acknowledge the use of the *Cryptococcus* gene deletion collection that was generated by the Madhani Laboratory at University of San Francisco and funded by the NIH funding (R01AI100272). We also acknowledge use of the *C. neoformans* genome sequences at the FungiDB database (http://fungidb.org/fungidb/). This study is supported by NIH grants AI123315 and AI155647 to C.X., and AI176922 to K.N. A.C. was supported in part by the National Institutes of Health (NIH) grants AI052733, AI15207, AI171093-01, and HL059842. The Xue lab is also supported by NIH grants AI169769 and AI141368.

## Author contributions

Y.W. and C.X. led the project, designed experiments, interpretated data and wrote the manuscript, with input from all authors. Y.W. performed many experiments and prepared figures. R.T. performed inositol uptake experiments and analyzed protein structural models. M.W. analyzed capsule structures, prepared figures and edited the manuscript. K.J. and K.N. provided the GWAS data of clinical isolates and edited the manuscript. Y.Z. performed animal experiments. Y.G. developed the protein structural models. K.N., A.C., and C.X. edited the manuscript and provide resources.

## Declaration of interestets

The authors declare no competing interests.

## Supplemental figures

**S1 Fig. Phenotypic analysis of group II *ITRs.* (a)** Cultures were grown overnight in YPD and diluted to an optical density at 600 nm (OD600) of 2.0. Tenfold serial dilutions were made in dH2O, and 5 ul of each was plated. The plates were grown for 2 days at 30°C and 37 °C. **(b)** Melanin detection assay. Color development by *itr4Δ, itr5Δ,* and *itr6Δ* single mutants and their triple mutants (*itr4Δ itr5Δ itr6Δ*) and wildtype strains was evaluated after cultures were grown on Niger Seed plate for 48 h at 30 °C and 37 °C. **(c)** Cultures were grown overnight in YPD and diluted to an OD600 of 2.0. Serial dilutions and spotting assays were made, and the plates were grown for 2 days at 30°C and 37 °C for different stress and inhibitor conditions. Wildtype and mutant strains are indicated on the left and the conditions at the top.

**S2 Fig.** Results from the virtual alanine mutation scan from the protein builder tool in Molecular Operation Environment (MOE). Receptor surface maps within 4.5Å of the ligand, shown as line diagrams, represent the shapes of the binding pockets in the wild-type and mutated forms of the protein. Red lines represent a negatively charged environment, blue lines represent a positively charged environment, and white lines represent neutral environments. The Q388A mutation had a significantly larger impact on the size of the binding pocket shape and volume than the Q389A mutation. Ligand interaction diagrams represent the non-covalent interactions between the ligand and receptor. Potential energy calculations computed did not account for hydration energies and may not fully capture the reality of the living cell.

**S3 Fig. Structural characterization of GXM polymers**. **(a)** SEC-MALS analysis of EPS from KN99, *itr4Δ,* and *ITR4^Q388,389A^* strains in glucose growth conditions. **(b)** 1D [^1^H] NMR analysis of EPS from all 3 strains. **(c)** Characterizing GXM motif expression and GXM O-acetylation.

**S4 Fig. Itr4 is required for fungal virulence in *in vitro* macrophage assay and *in vivo* animal models of systemic cryptococcosis.** (A) Macrophage killing assays were performed to assess the survival of fungal strains following co-culture with macrophages 4 and 24 h. Representative spotting assay images show fungal growth after macrophage interaction. (B) Quantification of fungal survival is presented as the surviving CFU relative to the control. Data represent individual biological replicates with bars showing the mean ± SEM. “*”, P < 0.05, “**”, P < 0.01; “ns”, not significant (determined by One-way ANOVA nonparametric test for multiple comparisons). (C) Expression profile of cytokine genes and genes involved in macrophage polarization was examined by qRT-PCR in mouse lung at day 7 after infection. Samples from naïve mice were analyzed as a control. Relative expression levels of indicated genes were calculated using the ΔΔCt method and normalized to GAPDH gene. Data are presented as individual biological replicates with bars representing the mean ± SEM. (D) Histological examination of lung sections. The lung sections were fixed and stained with H&E and GMS. Fungal cells appear dark brown or black with GMS staining. Yeast cells are highlighted with black arrow. (E) End point fungal burdens in the lungs, brains, spleens of animals infected with indicated strains in a murine intranasal inhalation model of systemic cryptococcosis. Each symbol represents one mouse. Bars represent the mean values +/- standard errors of the means. “****”, P *<* 0.0001; “*”, P *<* 0.001 “**”, P *<* 0.01; “*”, P *<* 0.05 (determined by One-way ANOVA nonparametric test for multiple comparisons).

**S5 Fig. Proposed model for the role of Itr4 in coordinating carbon utilization and fungal virulence.** Itr4 as an inositol receptor that is required to coordinate inositol and glucose metabolism by regulating ITRs and glucose sensor Hxs1. Itr4 regulates fungal pathogenesis in part through its regulation of capsule development. Itr4 serves as a critical sensor in the process of evolutionary specialization in this model.

## References

1. Rajasingham R, Smith RM, Park BJ, Jarvis JN, Govender NP, Chiller TM, et al. Global burden of disease of HIV-associated cryptococcal meningitis: an updated analysis. Lancet Infect Dis. 2017;17(8):873–81. Epub 20170505. doi: 10.1016/S1473-3099(17)30243-8. PubMed PMID: 28483415; PubMed Central PMCID: PMCPMC5818156.

2. Frases S, Pontes B, Nimrichter L, Viana NB, Rodrigues ML, Casadevall A. Capsule of *Cryptococcus neoformans* grows by enlargement of polysaccharide molecules. Proc Natl Acad Sci U S A. 2009;106(4):1228–33. Epub 20090121. doi: 10.1073/pnas.0808995106. PubMed PMID: 19164571; PubMed Central PMCID: PMCPMC2633523.

3. van der Horst CM, Saag MS, Cloud GA, Hamill RJ, Graybill JR, Sobel JD, et al. Treatment of cryptococcal meningitis associated with the acquired immunodeficiency syndrome. National Institute of Allergy and Infectious Diseases Mycoses Study Group and AIDS Clinical Trials Group. N Engl J Med. 1997;337(1):15–21. doi: 10.1056/NEJM199707033370103. PubMed PMID: 9203426.

4. Powderly WG. Cryptococcal Meningitis in HIV-Infected Patients. Curr Infect Dis Rep. 2000;2(4):352–7. doi: 10.1007/s11908-000-0015-y. PubMed PMID: 11095877.

5. Steenbergen JN, Nosanchuk JD, Malliaris SD, Casadevall A. *Cryptococcus neoformans* virulence is enhanced after growth in the genetically malleable host Dictyostelium discoideum. Infect Immun. 2003;71(9):4862–72. doi: 10.1128/IAI.71.9.4862-4872.2003. PubMed PMID: 12933827; PubMed Central PMCID: PMCPMC187309.

6. Cordero RJ, Frases S, Guimaraes AJ, Rivera J, Casadevall A. Evidence for branching in cryptococcal capsular polysaccharides and consequences on its biological activity. Mol Microbiol. 2011;79(4):1101–17. Epub 20110105. doi: 10.1111/j.1365-2958.2010.07511.x. PubMed PMID: 21208301; PubMed Central PMCID: PMCPMC3035750.

7. Cox GM, Mukherjee J, Cole GT, Casadevall A, Perfect JR. Urease as a virulence factor in experimental cryptococcosis. Infect Immun. 2000;68(2):443–8. doi: 10.1128/IAI.68.2.443-448.2000. PubMed PMID: 10639402; PubMed Central PMCID: PMCPMC97161.

8. Alspaugh JA, Perfect JR, Heitman J. *Cryptococcus neoformans* mating and virulence are regulated by the G-protein alpha subunit GPA1 and cAMP. Genes Dev. 1997;11(23):3206–17. PubMed PMID: 9389652.

9. Xue C, Tada Y, Dong X, Heitman J. The human fungal pathogen *Cryptococcus* can complete its sexual cycle during a pathogenic association with plants. Cell Host Microbe. 2007;1(4):263–73. doi: 10.1016/j.chom.2007.05.005. PubMed PMID: 18005707.

10. Pisani F, Livermore T, Rose G, Chubb JR, Gaspari M, Saiardi A. Analysis of Dictyostelium discoideum inositol pyrophosphate metabolism by gel electrophoresis. PLoS One. 2014;9(1). PubMed Central PMCID: PMCPMC3887064.

11. Xue C, Liu T, Chen L, Li W, Liu I, Kronstad JW, et al. Role of an expanded inositol transporter repertoire in *Cryptococcus neoformans* sexual reproduction and virulence. mBio. 2010;1(1). Epub 20100518. doi: 10.1128/mBio.00084-10. PubMed PMID: 20689743; PubMed Central PMCID: PMCPMC2912663.

12. Xue C. *Cryptococcus* and beyond--inositol utilization and its implications for the emergence of fungal virulence. PLoS Pathog. 2012;8(9). PubMed Central PMCID: PMCPMC3441655.

13. Liu TB, Kim JC, Wang Y, Toffaletti DL, Eugenin E, Perfect JR, et al. Brain inositol is a novel stimulator for promoting *Cryptococcus* penetration of the blood-brain barrier. PLoS Pathog. 2013;9(4):e1003247. Epub 20130404. doi: 10.1371/journal.ppat.1003247. PubMed PMID: 23592982; PubMed Central PMCID: PMCPMC3617100.

14. Donahue TF, Henry SA. Inositol Mutants of SACCHAROMYCES CEREVISIAE: Mapping the ino1 Locus and Characterizing Alleles of the ino1, ino2 and ino4 Loci. Genetics. 1981;98(3):491–503. doi: 10.1093/genetics/98.3.491. PubMed PMID: 17249096; PubMed Central PMCID: PMCPMC1214455.

15. Reynolds TB. Strategies for acquiring the phospholipid metabolite inositol in pathogenic bacteria, fungi and protozoa: making it and taking it. Microbiology (Reading). 2009;155(Pt 5):1386–96. Epub 20090421. doi: 10.1099/mic.0.025718-0. PubMed PMID: 19383710; PubMed Central PMCID: PMCPMC2889408.

16. Lai K, Bolognese CP, Swift S, McGraw P. Regulation of inositol transport in *Saccharomyces cerevisiae* involves inositol-induced changes in permease stability and endocytic degradation in the vacuole. J Biol Chem. 1995;270(6):2525–34. doi: 10.1074/jbc.270.6.2525. PubMed PMID: 7852314.

17. Lai K, McGraw P. Dual control of inositol transport in *Saccharomyces cerevisiae* by irreversible inactivation of permease and regulation of permease synthesis by INO2, INO4, and OPI1. J Biol Chem. 1994;269(3):2245–51. PubMed PMID: 8294482.

18. Robinson KS, Lai K, Cannon TA, McGraw P. Inositol transport in *Saccharomyces cerevisiae* is regulated by transcriptional and degradative endocytic mechanisms during the growth cycle that are distinct from inositol-induced regulation. Mol Biol Cell. 1996;7(1):81–9. doi: 10.1091/mbc.7.1.81. PubMed PMID: 8741841; PubMed Central PMCID: PMCPMC278614.

19. Chen YL, Kauffman S, Reynolds TB. *Candida albicans* uses multiple mechanisms to acquire the essential metabolite inositol during infection. Infect Immun. 2008;76(6):2793–801. Epub 20080211. doi: 10.1128/IAI.01514-07. PubMed PMID: 18268031; PubMed Central PMCID: PMCPMC2423082.

20. Jin JH, Seyfang A. High-affinity myo-inositol transport in *Candida albicans*: substrate specificity and pharmacology. Microbiology (Reading). 2003;149(Pt 12):3371–81. doi: 10.1099/mic.0.26644-0. PubMed PMID: 14663071.

21. Wang Y, Liu TB, Delmas G, Park S, Perlin D, Xue C. Two major inositol transporters and their role in cryptococcal virulence. Eukaryot Cell 2011;10(5):618–28. doi: 10.1128/EC.00327-10. PubMed Central PMCID: PMCPMC3127654.

22. Boulware DR, Meya DB, Muzoora C, Rolfes MA, Huppler Hullsiek K, Musubire A, et al. Timing of antiretroviral therapy after diagnosis of cryptococcal meningitis. N Engl J Med. 2014;370(26):2487-98. doi: 10.1056/NEJMoa1312884. PubMed PMID: 24963568; PubMed Central PMCID: PMCPMC4127879.

23. Gerstein AC, Jackson KM, McDonald TR, Wang Y, Lueck BD, Bohjanen S, et al. Identification of Pathogen Genomic Differences That Impact Human Immune Response and Disease during *Cryptococcus neoformans* Infection. mBio. 2019;10(4). Epub 20190716. doi: 10.1128/mBio.01440-19. PubMed PMID: 31311883; PubMed Central PMCID: PMCPMC6635531.

24. Jackson KM, Kono TJY, Betancourt JA-O, Wang YA-OX, Kabbale KD, Ding M, et al. Single nucleotide polymorphisms are associated with strain-specific virulence differences among clinical isolates of *Cryptococcus neoformans*. Nat Commun. 2024;15(1).

25. Wang Y, Wear M, Kohli G, Vij R, Giamberardino C, Shah A, et al. Inositol Metabolism Regulates Capsule Structure and Virulence in the Human Pathogen *Cryptococcus neoformans*. mBio. 2021;12(6):e0279021. Epub 20211102. doi: 10.1128/mBio.02790-21. PubMed PMID: 34724824; PubMed Central PMCID: PMCPMC8561382.

26. Bahn YS, Xue C, Idnurm A, Rutherford JC, Heitman J, Cardenas ME. Sensing the environment: lessons from fungi. Nat Rev Microbiol. 2007;5(1):57–69. doi: 10.1038/nrmicro1578. PubMed PMID: 17170747.

27. Ding M, Jackson KM, Harris-Gordon M, Dileepan T, Meya DB, Nielsen K. The hypervirulent Type-1/Type-17 phenotype of *Cryptococcus neoformans* clinical isolates is specific to A/J mice. Infect Immun. 2025;93(4):e0058524. Epub 20250303. doi: 10.1128/iai.00585-24. PubMed PMID: 40029251; PubMed Central PMCID: PMCPMC11977316.

28. Wieczorke R, Krampe S, Weierstall T, Freidel K, Hollenberg CP, Boles E. Concurrent knock-out of at least 20 transporter genes is required to block uptake of hexoses in *Saccharomyces cerevisiae*. FEBS Lett. 1999;464(3):123–8. doi: 10.1016/s0014-5793(99)01698-1. PubMed PMID: 10618490.

29. Liu TB, Wang Y, Baker GM, Fahmy H, Jiang L, Xue C. The glucose sensor-like protein Hxs1 is a high-affinity glucose transporter and required for virulence in *Cryptococcus neoformans*. PLoS One. 2013;8(5):e64239. Epub 20130514. doi: 10.1371/journal.pone.0064239. PubMed PMID: 23691177; PubMed Central PMCID: PMCPMC3653957.

30. Janbon G, Ormerod KL, Paulet D, Byrnes EJ, 3rd, Yadav V, Chatterjee G, et al. Analysis of the genome and transcriptome of *Cryptococcus neoformans var. grubii* reveals complex RNA expression and microevolution leading to virulence attenuation. PLoS Genet. 2014;10(4). PubMed Central PMCID: PMCPMC3990503.

31. Treitel MA, Kuchin S Fau - Carlson M, Carlson M. Snf1 protein kinase regulates phosphorylation of the Mig1 repressor in *Saccharomyces cerevisiae*. Mol Cell Biol. 1998;18(11):6273–80. doi: 10.1128/MCB.18.11.6273. PubMed Central PMCID: PMCPMC109214.

32. Caza M, Hu G, Price M, Perfect JR, Kronstad JW. The Zinc Finger Protein Mig1 Regulates Mitochondrial Function and Azole Drug Susceptibility in the Pathogenic Fungus *Cryptococcus neoformans*. LID - 10.1128/mSphere.00080-15 [doi] LID - e00080-15. mSphere. 2016;1(1). PubMed Central PMCID: PMCPMC4863601.

33. Kayikci Ö, Nielsen J. Glucose repression in *Saccharomyces cerevisiae*. LID - fov068 [pii] LID - 10.1093/femsyr/fov068 [doi]. FEMS Yeast Res 2015;15(6):fov068. doi: 10.1093/femsyr/fov068. PubMed Central PMCID: PMCPMC4629793.

34. Gancedo JM. Yeast carbon catabolite repression. Microbiol Mol Biol Rev. 1998;62(2):334–61. doi: 10.1128/MMBR.62.2.334-361.; PubMed Central PMCID: PMCPMC98918.

35. Kaniak A, Xue Z, Macool D, Kim JH, Johnston M. Regulatory network connecting two glucose signal transduction pathways in *Saccharomyces cerevisiae*. Eukaryot Cell. 2004;3(1):221–31. doi: 10.1128/EC.3.1.221-231.2004. PubMed PMID: 14871952; PubMed Central PMCID: PMCPMC329515.

36. Hedbacker K, Carlson M. SNF1/AMPK pathways in yeast. Front Biosci. 2008;13:2408–20. PubMed Central PMCID: PMCPMC2685184.

37. Jumper J, Evans R, Pritzel A, Green T, Figurnov M, Ronneberger O, et al. Highly accurate protein structure prediction with AlphaFold. Nature. 2021;596(7873):583–9. Epub 20210715. doi: 10.1038/s41586-021-03819-2. PubMed PMID: 34265844; PubMed Central PMCID: PMCPMC8371605.

38. Moriya H, Johnston M. Glucose sensing and signaling in *Saccharomyces cerevisiae* through the Rgt2 glucose sensor and casein kinase I. Proc Natl Acad Sci U S A. 2004;101(6):1572–7. Epub 20040130. doi: 10.1073/pnas.0305901101. PubMed PMID: 14755054; PubMed Central PMCID: PMCPMC341776.

39. Frases S, Nimrichter L, Viana NB, Nakouzi A, Casadevall A. *Cryptococcus neoformans* capsular polysaccharide and exopolysaccharide fractions manifest physical, chemical, and antigenic differences. Eukaryot Cell. 2008;7(2):319–27. Epub 20071221. doi: 10.1128/EC.00378-07. PubMed PMID: 18156290; PubMed Central PMCID: PMCPMC2238165.

40. Alvarez M, Casadevall A. Phagosome extrusion and host-cell survival after Cryptococcus neoformans phagocytosis by macrophages. Curr Biol. 2006;16(21):2161–5. doi: 10.1016/j.cub.2006.09.061. PubMed PMID: 17084702.

41. Ma H, Croudace JE, Lammas DA, May RC. Expulsion of live pathogenic yeast by macrophages. Curr Biol. 2006;16(21):2156–60. doi: 10.1016/j.cub.2006.09.032. PubMed PMID: 17084701.

42. Fu MS, Liporagi-Lopes LC, Dos Santos SRJ, Tenor JL, Perfect JR, Cuomo CA-OX, et al. Amoeba Predation of Cryptococcus neoformans Results in Pleiotropic Changes to Traits Associated with Virulence. LID - 10.1128/mBio.00567-21 [doi] LID - e00567-21. mBio. 2021;12(2). PubMed Central PMCID: PMCPMC8092252.

43. Steenbergen JN, Shuman Ha Fau - Casadevall A, Casadevall A. *Cryptococcus neoformans* interactions with amoebae suggest an explanation for its virulence and intracellular pathogenic strategy in macrophages. Proc Natl Acad Sci U S A. 2001;98(26). PubMed Central PMCID: PMCPMC65014.

44. Levitz SM. Does amoeboid reasoning explain the evolution and maintenance of virulence factors in *Cryptococcus neoformans*? Proc Natl Acad Sci, U S A. 2001;98(26). PubMed Central PMCID: PMCPMC64930.

45. Manske C, Schell U, Hilbi H. Metabolism of myo-Inositol by *Legionella pneumophila* Promotes Infection of Amoebae and Macrophages. Appl Environ Microbiol. 2016;82(16):5000–14. Epub 20160729. doi: 10.1128/AEM.01018-16. PubMed PMID: 27287324; PubMed Central PMCID: PMCPMC4968532.

46. Yu CH, Sephton-Clark P, Tenor JL, Toffaletti DL, Giamberardino C, Haverkamp M, et al. Gene Expression of Diverse *Cryptococcus* Isolates during Infection of the Human Central Nervous System. mBio. 2021;12(6):e0231321. Epub 20211102. doi: 10.1128/mBio.02313-21. PubMed PMID: 34724829; PubMed Central PMCID: PMCPMC8561399.

47. Lorenz MC, Heitman J. The MEP2 ammonium permease regulates pseudohyphal differentiation in Saccharomyces cerevisiae. EMBO J. 1998;17(5):1236–47. doi: 10.1093/emboj/17.5.1236. PubMed PMID: 9482721; PubMed Central PMCID: PMCPMC1170472.

48. Rutherford JC, Chua G, Hughes T, Cardenas ME, Heitman J. A Mep2-dependent transcriptional profile links permease function to gene expression during pseudohyphal growth in *Saccharomyces cerevisiae*. Mol Biol Cell. 2008;19(7):3028–39. Epub 20080423. doi: 10.1091/mbc.e08-01-0033. PubMed PMID: 18434596; PubMed Central PMCID: PMCPMC2441671.

49. van den Berg B, Chembath A, Jefferies D, Basle A, Khalid S, Rutherford JC. Structural basis for Mep2 ammonium transceptor activation by phosphorylation. Nat Commun. 2016;7(2041–1723 (Electronic)):11337. Epub 20160418. doi: 10.1038/ncomms11337. PubMed PMID: 27088325; PubMed Central PMCID: PMCPMC4852598.

50. Carlson M. Glucose repression in yeast. Curr Opin Microbiol. 1999;2(2):202–7. doi: 10.1016/S1369-5274(99)80035-6. PubMed PMID: 10322167.

51. Santangelo GM. Glucose signaling in *Saccharomyces cerevisiae*. Microbiol Mol Biol Rev. 2006;70(1):253–82. doi: 10.1128/MMBR.70.1.253-282.2006. PubMed PMID: 16524925; PubMed Central PMCID: PMCPMC1393250.

52. Doering TL. How sweet it is! Cell wall biogenesis and polysaccharide capsule formation in *Cryptococcus neoformans*. Annu Rev Microbiol. 2009;63:223–47.

53. Granger Dl Fau - Perfect JR, Perfect Jr Fau - Durack DT, Durack DT. Virulence of *Cryptococcus neoformans*. Regulation of capsule synthesis by carbon dioxide. J Clin Invest. 1985;76(2):508–16.

54. Xue C, Bahn YS, Cox GM, Heitman J. G protein-coupled receptor Gpr4 senses amino acids and activates the cAMP-PKA pathway in *Cryptococcus neoformans*. Mol Biol Cell. 2006;17(2):667–79. PubMed PMID: 16291861; PubMed Central PMCID: PMC1356578.

55. Bahn YS, Kojima K Fau - Cox GM, Cox Gm Fau - Heitman J, Heitman J. Specialization of the HOG pathway and its impact on differentiation and virulence of Cryptococcus neoformans. Mol Biol Cell. 2005;16(5):2285–300. doi: 10.1091/mbc.e04-11-0987. PubMed Central PMCID: PMCPMC1087235.

56. Davidson RC, Blankenship JR, Kraus PR, de Jesus Berrios M, Hull CM, D’Souza C, et al. A PCR-based strategy to generate integrative targeting alleles with large regions of homology. Microbiology (Reading). 2002;148(Pt 8):2607–15. doi: 10.1099/00221287-148-8-2607. PubMed PMID: 12177355.

57. Liu TB, Wang Y Fau - Stukes S, Stukes S Fau - Chen Q, Chen Q Fau - Casadevall A, Casadevall A Fau - Xue C, Xue C. The F-Box protein Fbp1 regulates sexual reproduction and virulence in *Cryptococcus neoformans*. Eukaryot Cell. 2011;10(6):791–802. doi: 10.1128/EC.00004-11. PubMed Central PMCID: PMCPMC3127668.

58. Hankins HM, Sere YY, Diab NS, Menon AK, Graham TR. Phosphatidylserine translocation at the yeast trans-Golgi network regulates protein sorting into exocytic vesicles. Mol Biol Cell. 2015;26(25):4674–85. Epub 20151014. doi: 10.1091/mbc.E15-07-0487. PubMed PMID: 26466678; PubMed Central PMCID: PMCPMC4678023.

59. Liu TB, Xue C. Fbp1-mediated ubiquitin-proteasome pathway controls Cryptococcus neoformans virulence by regulating fungal intracellular growth in macrophages. Infect Immun. 2014;82(2):557–68. Epub 20131118. doi: 10.1128/IAI.00994-13. PubMed PMID: 24478071; PubMed Central PMCID: PMCPMC3911387.

60. Yao H, Kim K, Duan M, Hayashi T, Guo M, Morgello S, et al. Cocaine hijacks sigma1 receptor to initiate induction of activated leukocyte cell adhesion molecule: implication for increased monocyte adhesion and migration in the CNS. J Neurosci. 2011;31(16):5942–55. doi: 10.1523/JNEUROSCI.5618-10.2011. PubMed PMID: 21508219; PubMed Central PMCID: PMCPMC3410749.

61. Jong A, Wu CH, Prasadarao NV, Kwon-Chung KJ, Chang YC, Ouyang Y, et al. Invasion of *Cryptococcus neoformans* into human brain microvascular endothelial cells requires protein kinase C-alpha activation. Cell Microbiol. 2008;10(9):1854–65. Epub 20080516. doi: 10.1111/j.1462-5822.2008.01172.x. PubMed PMID: 18489726; PubMed Central PMCID: PMCPMC2729555.

62. Kim JC, Crary B, Chang YC, Kwon-Chung KJ, Kim KJ. *Cryptococcus neoformans* activates RhoGTPase proteins followed by protein kinase C, focal adhesion kinase, and ezrin to promote traversal across the blood-brain barrier. J Biol Chem. 2012;287(43):36147–57. Epub 20120816. doi: 10.1074/jbc.M112.389676. PubMed PMID: 22898813; PubMed Central PMCID: PMCPMC3476282.

63. Anders S, Huber W. Differential expression analysis for sequence count data. Genome Biol. 2010;11(10):R106. Epub 20101027. doi: 10.1186/gb-2010-11-10-r106. PubMed PMID: 20979621; PubMed Central PMCID: PMCPMC3218662.

64. Masso-Silva J, Espinosa V, Liu TB, Wang Y, Xue C, Rivera A. The F-Box Protein Fbp1 Shapes the Immunogenic Potential of *Cryptococcus neoformans*. mBio. 2018;9(1). Epub 20180109. doi: 10.1128/mBio.01828-17. PubMed PMID: 29317510; PubMed Central PMCID: PMCPMC5760740.

65. Wang Y, Wang K, Masso-Silva JA, Rivera A, Xue C. A Heat-Killed *Cryptococcus* Mutant Strain Induces Host Protection against Multiple Invasive Mycoses in a Murine Vaccine Model. mBio. 2019;10(6). Epub 20191126. doi: 10.1128/mBio.02145-19. PubMed PMID: 31772051; PubMed Central PMCID: PMCPMC6879717.

